# Biphasic glycocalyx remodelling balances immune cell recruitment and tissue damage during influenza infection

**DOI:** 10.64898/2026.11.16.766997

**Authors:** Iashia Z Nabi, Ibrahim A Ahmed, Rebecca J Dodd, James E Parkinson, Adam Johnson, Amanda JL Ridley, Ziyun Zhang, Kirstin O Lowe, Catherine Park, Samie Dalgarno, Lucy J Le Maistre, Judith E Allen, Jin-Ping Li, Antony D Adamson, Tracy Hussell, Douglas P Dyer

## Abstract

The endothelial glycocalyx is a glycan-rich barrier lining blood vessels that regulates immune cell recruitment, whose role in viral infection is unknown. Using a murine influenza model, we reveal that the lung endothelial glycocalyx undergoes coordinated biphasic remodelling: early shedding followed by substantial accumulation during peak immune activation. Genetic deletion of heparanase, the enzyme driving early glycocalyx shedding, reduced immune cell lung infiltration and improved host recovery. Conversely, using a new endothelial-specific glycocalyx depletion mouse model, we demonstrate that this glycocalyx accumulation actively limits immune cell recruitment and prevents excessive tissue damage. Together, these findings reveal that coordinated endothelial and immune cell glycocalyx remodelling is an active and previously unrecognised regulator of the balance between antiviral immune cell recruitment and tissue protection.

## Introduction

Immune cell recruitment is an essential component of the immune response, facilitating co-ordinated anti-viral immunity and subsequent recovery of tissue homeostasis^1^. Respiratory viral infection is a global health burden responsible for approximately 6 million deaths annually. During influenza A Virus (IAV) infection recruited immune cells, that clear the virus, can also damage the lung tissue that they are trying to protect. When this balance between defence and damage is dysregulated, severe disease and death can result^2–7^. How this process is controlled remains incompletely understood. Here we identify the endothelial glycocalyx (eGC), the glycan-rich barrier lining blood vessels^8–10^, as an active and previously unrecognised regulator of immune cell recruitment. We reveal that the glycocalyx operates through a coordinated biphasic remodelling programme, during influenza infection.

Influenza A virus (IAV) infection induces substantial pulmonary inflammation and immune-mediated lung pathology, dependent on viral load and host immune responses^11,12^. Immune cell recruitment during acute lung infection plays a central role in both infection and resolution, but also aberrant immune responses leading to chronic lung damage and more severe disease^2–6^. Therefore, understanding the mechanisms that regulate immune cell recruitment to achieve a balance between antiviral immunity and tissue damage is a crucial challenge^2,13^.

The cell surface glycocalyx surrounds both endothelial and immune cells and is a key regulator of immune cell recruitment and function^8,10^. However, the mechanisms of glycocalyx function remain poorly understood, particularly in the context of viral infection^9,10^. Proteoglycans are the major building blocks of the glycocalyx and are key immunoregulatory components^14^. They are composed of membrane-associated protein cores (e.g. syndecan-1 (SDC-1)) with glycosaminoglycan (GAG) side chains which are either sulfated (e.g. heparan sulfate (HS)), or non-sulfated GAG (e.g hyaluronan (HA)). In blood vessels the eGC lines the luminal surface and forms a thick, hydrated, adhesion-resistant barrier structure. Here the eGC acts as a key regulator of vascular homeostasis, endothelial barrier integrity and immune cell trafficking^15^.

Furthermore, immune cells also have cell-specific glycocalyces whose function we are only just beginning to appreciate in their activation, migration and inflammatory function^16–22^.

Endothelial, and immune-cell glycocalyx degradation (shedding) is a major feature of acute inflammatory diseases including sepsis, acute respiratory distress syndrome (ARDS), COVID-19 and influenza infection^23–27^. Such glycocalyx degradation and shedding is implicated in endothelial activation, increased vascular permeability, oedema formation and enhanced immune cell adhesion^9,10,28^. Despite this, few studies have investigated the temporal regulation, spatial localisation and function of specific pulmonary glycocalyx-associated components such as HS, HA and SDC-1 during viral lung infection.

Here we use a murine model of IAV infection to demonstrate that eGC components showed biphasic early shedding followed by substantial accumulation later in the antiviral immune response. To determine the function of eGC accumulation we developed a new mouse model to cell-specifically deplete a key glycocalyx synthesis gene (*B4Galt7*^29,30^). When this enzyme was targeted in endothelial cells, eGC barrier accumulation was ablated, resulting in increased immune cell recruitment and exacerbated tissue damage. Furthermore, genetic removal of the enzyme controlling early glycocalyx shedding (i.e. heparanase^31^) resulted in diminished immune cell recruitment and enhanced host recovery. Therefore, our study reveals that IAV infection induces coordinated eGC remodelling, which is closely linked to the inflammatory response, crucially balancing immune cell recruitment against excessive tissue damage.

## Results

### The eGC is shed early in the immune response to influenza infection

Shedding of the eGC is associated with immune cell recruitment and tissue pathology in a range of disease contexts^9,10,25,28^. Most of this evidence consists of increased levels of the glycocalyx components, e.g. the HS side chains from proteoglycans (Fig. 1A), in the serum of patients with a range of diseases including acute respiratory distress syndromes (ARDS)^24^. Classically, shed glycocalyx is attributed to release from endothelial cells. To definitively address this, we firstly used light sheet microscopy to reveal that the cleared mouse lung has an extensive overlap of CD31 (vascular marker) and HS (glycocalyx component) expression (Fig. 1B). This was validated using immunofluorescent staining of mouse lung sections with antibodies to VWF^+^ endothelial cells and HS (Fig. 1C). Closer inspection of this vascular HS staining relative to the endothelial cells lining the vasculature (DAPI^+^) depicted some HS positive staining on the basal endothelium (from the basement membrane) with the majority located on the apical endothelium (eGC) (Fig. 1D). Mouse lung endothelial vessels were also positive for another eGC component HA (Supp. Fig. 1A). These techniques confirmed the widespread presence of the eGC within the mouse lung vasculature.

**Figure 1.**
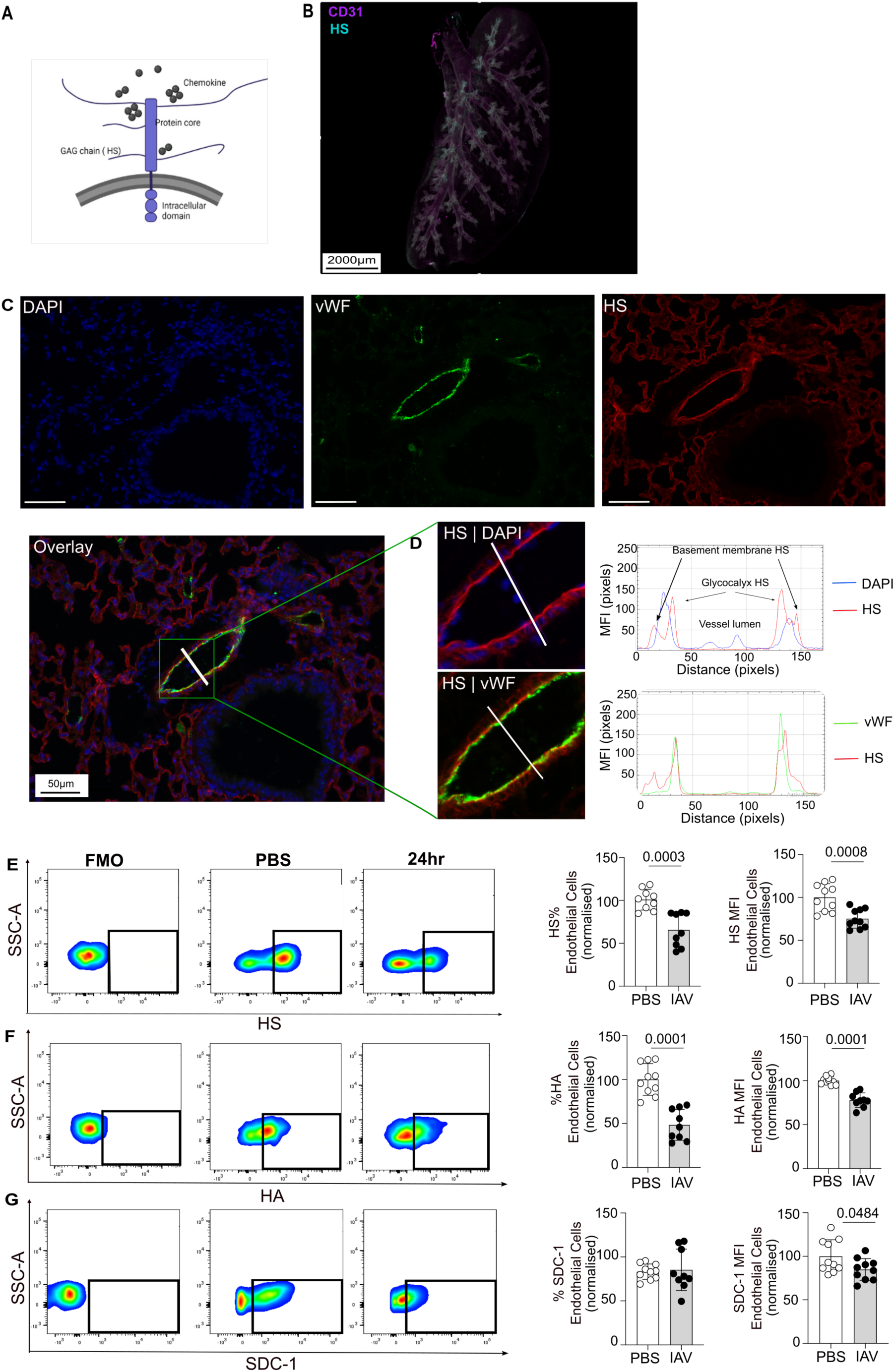
The pulmonary endothelial glycocalyx is shed early following IAV infection. (A) Schematic of an example proteoglycan structure (SDC-1 with HS), which are key glycocalyx components. (B) Whole organ imaging of naïve mouse lung that has been cleared and stained for blood vessels (CD31; magenta) and HS (cyan). Scale bar is 2000µm. (C) Immunofluorescence of FFPE naïve mouse lung sections stained for endothelial cells (VWF; green), HS (red) and DAPI (blue) with merged overlay. Scale bars represent 50 µm (D) Relative localisation of vWF and HS around endothelial cells of a naïve mouse pulmonary vessel with line profile of MFI. Flow cytometry staining plots and associated quantification of endothelial (CD45^-^CD31^+^) levels of (E) HS, (F) HA and (G) SDC-1, normalised relative to levels in PBS controls. (E–G) each point represents an individual mouse (n=8-10), data are plotted as mean ± SD. E-G were analysed by unpaired t-test with p values indicated.

We next utilised flow cytometry to specifically analyse the glycocalyx content on vascular endothelial cells (gating in Supp. Fig. 1B) early (24 hours) after infection, when recruitment of immune cells to the lung was first observed (Supp. Fig. 1C-D). At 24 hours post infection we observed reduced endothelial levels of the glycocalyx components HS (Fig. 1E), HA (Fig. 1F) and SDC-1 (Fig. 1G). These data support the hypothesis that the pulmonary vasculature contribute to the shed glycocalyx components detected in serum during viral infection.

### The eGC accumulates during the transition between the late innate and adaptive phase of the murine viral infection time course

To broaden this classical “shedding” concept, we analysed the eGC across the time course of influenza A infection. Firstly, we analysed a mouse lung single-cell dataset across the infection time course^32,33^ from a glycocalyx perspective (Supp. Fig 2A). Given the initial shedding of the glycocalyx from endothelial cells observed 24 hrs after IAV infection, we plotted the temporal expression of genes encoding glycocalyx remodelling enzymes in different endothelial subsets. Specifically heparanase (*Hpse*) (which cleaves HS chains directly^31^), *Sulf 1* and *Sulf 2* (which remodel HS sulfation^34,35^) and *Mmp2* (which cleaves proteoglycan cores^36^) (Fig. 2A) along with other *Mmp’s* (Supp. Fig. 2B). These data revealed dynamic glycocalyx remodelling gene expression across the IAV infection time course, specific to individual endothelial cell populations. Furthermore, we reveal that VWF^+^ endothelial subsets, which facilitate immune cell recruitment during inflammation^1,37^, have the highest expression of glycocalyx remodelling genes. For example, *Hpse* expression in VWF^+^ endothelial cells initially decreased post infection and then peaked around 7 days post infection (dpi). This time point coincides with peak expression of *Sulf1, Sulf2, Mmp2, Mmp3* and *Mmp14* in the VWF^+^ endothelial subset (Fig. 2A and Supp. Fig. 2B). Lymphatic endothelial cells (Prox1^+^)^38^ also expressed glycocalyx remodelling genes, with *Sulf2* and *Mmp14* being dominant in these cells. This reflects glycocalyx remodelling that is both time-dependent and specific to endothelial cell subsets, mirroring discrete phases of the immune response.

**Figure 2.**
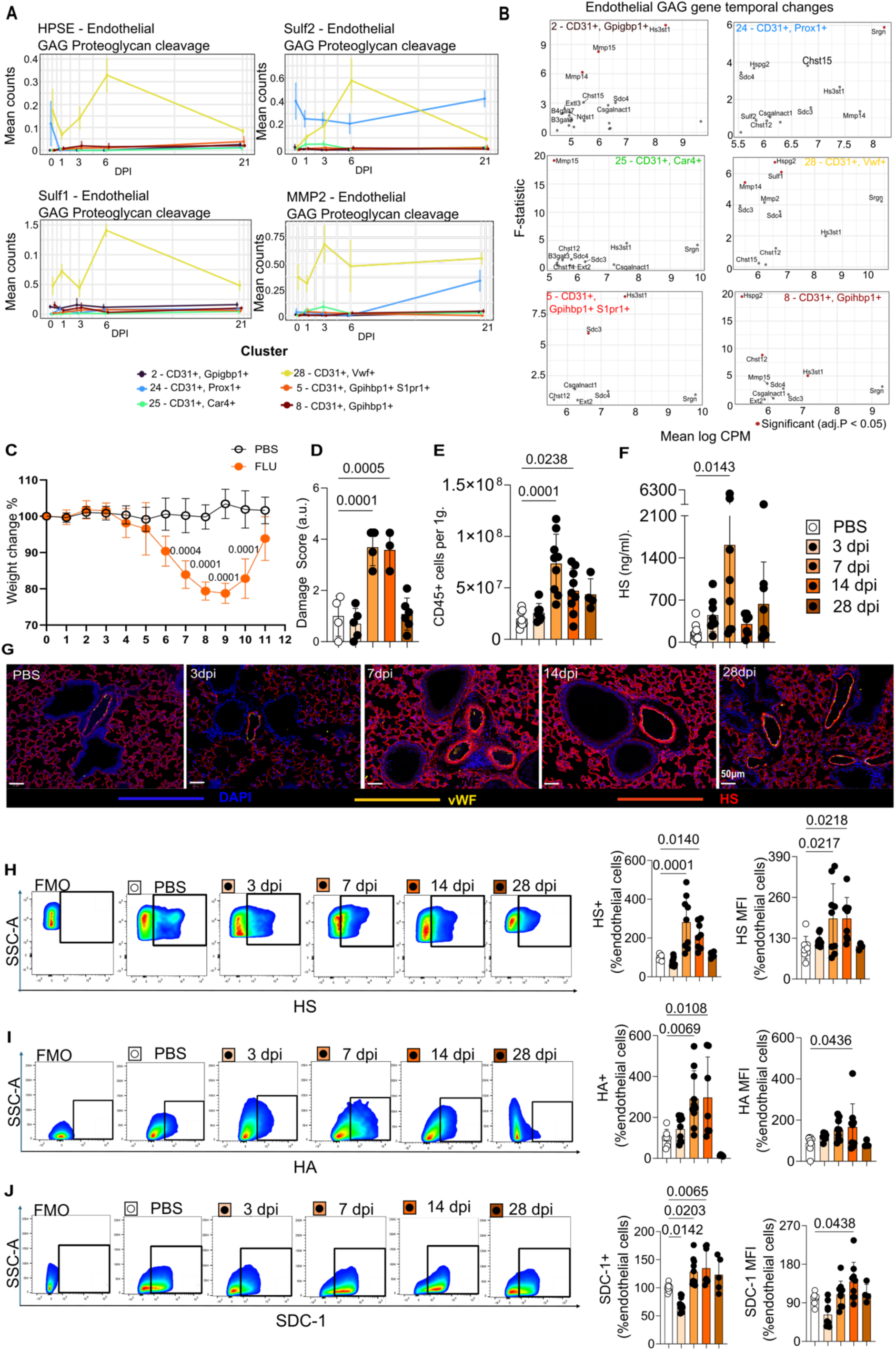
The pulmonary endothelial glycocalyx barrier accumulates 7 dpi. (A) Expression analysis of genes transcribing glycocalyx remodelling enzymes HPSE, SULF1, SULF2 and MMP-2 in different endothelial subsets across the IAV infection time course. (B) Differential expression analysis of the most dynamic GAG synthesis genes across the influenza infection time course in endothelial subsets. (C) Weight change in WT control and influenza infected mice up to day 11 post infection. (D) Damage scoring of mouse lung sections, (E) Flow-cytometry based quantification of immune cells (CD45^+^) in mouse lungs, (F) ELISA based quantification of HS serum levels, (G) Representative immunofluorescence images of FFPE tissue sections from WT mice for cell nucleus (DAPI; blue), vWF (yellow)and HS (red); day 0-28 post influenza infection. Scale bars represent 50 µm. Flow cytometry staining and associated analysis of endothelial cell (CD45^-^CD31^+^) positivity and MFI for (H) HS, (I), HA and (J) SDC-1 in WT mice days 0-28 post influenza infection normalised to PBS controls. Each point represents an individual mouse, (D; n=3-5), (C, E-F and H-J, n=6-10), data are plotted as mean ± SD. C was analysed by two-way ANOVA and D-F and H-J were analysed by one-way ANOVA with p values indicated.

As a result of these temporal changes, we next analysed the dataset for genes involved in GAG regulation (synthesis and degradation) that changed across the IAV infection time course (F-statistic) in the different endothelial subsets (Fig. 2B). Again VWF^+^ endothelial cells showed the biggest changes in gene expression across the IAV infection time course, e.g. the protein core encoding *Sdc3 and Sdc4*, GAG synthesis encoding *Hs3st1* and the glycocalyx remodelling genes *Sulf1* and *Mmp*s. Gene expression changes were also observed in GAG synthesis and degradation genes across the endothelial cell subsets. To further understand the underlying transcriptional response, we then analysed expression of grouped protein core and GAG synthesis genes in the individual endothelial subsets (Supp. Fig 2C), across all identified cell types (Supp. Fig. 2D) and also individual GAG synthesis gene expression in endothelial subsets (Supp. Fig. 2E) across the IAV infection time course. Together, these transcriptional data predict dynamic, cell-specific remodelling of the eGC across the IAV infection time course.

To analyse the functional outcome of these temporal gene expression patterns we performed a murine IAV model of infection, lung injury and resolution (Supp. Fig. 3A) to probe the function, and remodelling of, the eGC. This model produced weight loss (Fig. 2C), tissue damage (Fig. 2D and Supp. Fig. 3B-C), immune cell recruitment (Fig. 2E and Supp. Fig. 3D-E) and HS glycocalyx shedding into the serum (Fig. 2F). All these parameters peaked approximately 7 days after infection and resolved towards baseline levels 28 days after infection.

We stained lung tissues sections, taken from 0, 3, 7, 14 and 28 dpi for vWF (endothelial marker) and HS GAG (Fig 2G). These images revealed that instead of returning to baseline following initial shedding at 24 hours post infection, the endothelial HS at days 7 and 14 dpi unexpectedly accumulates to levels higher than those at rest. To confirm this observation, we used flow cytometry-based analysis of the eGC content across the time course of infection. These data again demonstrated significantly higher levels of HS (Fig. 2H) and HA (Fig. 2I) GAGs in addition to the proteoglycan core SDC-1 (Fig. 2J) on endothelial cells, at 7 and 14 dpi, compared to uninfected controls. Together these data reveal, for the first time, that the eGC is significantly increased during the transition from the late innate to adaptive phase (7-14 dpi), of the IAV infection time course. Crucially the transition from late innate to the adaptive phase is when virus is cleared and the immune response is dampened to prevent excessive inflammation and tissue damage. As such this stage of viral infection is key in the balance between viral clearance and tissue damage, which are both regulated by immune cell recruitment^2^.

### Accumulation of the eGC regulates immune cell recruitment and tissue damage following viral infection

Having demonstrated that the eGC accumulates during the key transition from the late innate to adaptive phase of the viral immune response, we hypothesised that this accumulation limits immune cell recruitment, which drives tissue damage following viral clearance. To test this hypothesis, we generated a new mouse model introducing loxP sites around the gene for *B4Galt7* (Fig. 3A and Supp. Fig. 4A). This enzyme facilitates attachment of the sulfated GAGs HS and CS onto protein cores^29^, therefore its removal should significantly reduce the GAG content of the glycocalyx in a cell specific manner. This mouse was crossed to the inducible Cadherin-5 (Cdh5) endothelial cell-specific Cre mouse line. Subsequently we confirmed recombination and gene deletion in endothelial (CD45^-^CD31^+^), but not immune (CD45^+^), cells following tamoxifen treatment (Supp. Fig. 4B-E). To determine whether removal of *B4Galt7* impacted the eGC, we first stained tissues from IAV-infected WT and endothelial specific knockout (KO) mice for VWF and HS. This approach visually demonstrated reduction of vascular HS levels (Fig. 3B) in mice with *B4Galt7* deficient endothelial cells. This result was reinforced by flow cytometric analysis of endothelial cell HS (Fig. 3C), confirming that the virally induced accumulation of HS observed at day 7 post infection, was prevented when endothelial cells are deficient in *B4Galt7*.

**Figure 3.**
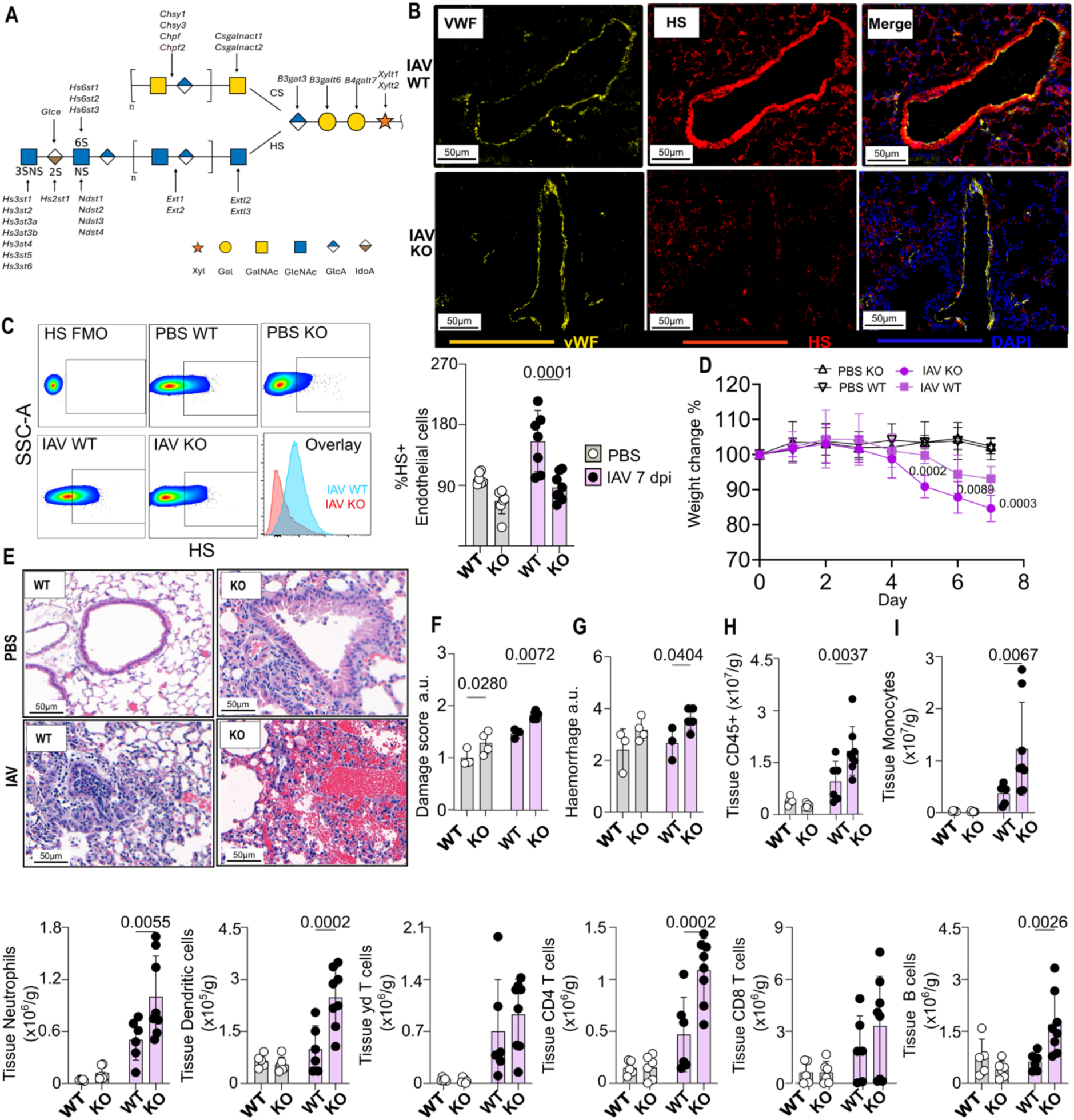
The eGC barrier regulates the balance between immune cell recruitment and tissue damage following influenza infection. (A) Schematic of genes required for synthesis and sulfation of HS and CS GAGs. (B) Representative FFPE Immunofluorescence of lung tissue section stained for cell nucleus (DAPI; blue), endothelial cells (VWF; yellow) and HS (red) and (C) flow cytometry analysis and quantification of endothelial (CD45^-^CD31^+^) HS levels, (D) Quantification of weight, (E) Haematoxylin and eosin (H&E) staining of mouse lung sections and associated (F) overall tissue damage score and (G) haemorrhage score; in wild type (Cdh5-Cre/ERT2^-/-^x B4Galt7^fl/fl^) and endothelial GAG KO (Cdh5-Cre/ERT2^+/-^ x B4Galt7^fl/fl^) mice 7 days after control or influenza infection. Flow cytometry-based quantification of mouse lung tissue for (H) overall immune cells (CD45^+^) and (I) individual immune cell populations. Each dot is an individual mouse (F-G, J; n=3-5), (D, and H-I, n=6-10), data are plotted as mean ± SD. Data from C-I were analysed by two-way ANOVA with p values indicated.

Having developed this new tool, we next applied it to probe the role of eGC accumulation during the IAV immune response. First, we observed that weight loss was significantly exacerbated in the *B4Galt7* endothelial cell KO mice 5-7 dpi (Fig. 3D), evidence of a worsened host response to viral infection. We applied our histological tissue damage analysis to these same mice and confirmed that tissue damage was significantly worsened in the absence of the HS glycocalyx barrier (Fig. 3E-F and Supp. Fig 4F). The overall worsened score was driven by increased levels of haemorrhage following viral infection (Fig. 3G), in endothelial cell specific *B4Galt7* depleted mice. Furthermore, these mice also had increased numbers of CD45^+^ immune cells in their lung tissue, compared to IAV infected WT controls (Fig. 3H). Specifically, monocytes, neutrophils, dendritic cells (DCs) and CD4 T cells were increased compared to wild type controls (Fig. 3I). These immune cells are typically present at this infection time point in wild type mice (Supp. Fig 3); demonstrating a wide-spread function for this accumulated eGC in controlling immune cell recruitment in this model.

Interestingly, we also saw significantly higher levels of B cells at 7 dpi in the *B4Galt7* KO mice (Fig. 3I), which are not significantly increased at this time point in WT mice (Supp. Fig 3); suggesting that the nature of the immune response is also being controlled by the eGC. Together these data demonstrated that the increased eGC following viral infection is a key regulator in balancing immune cell recruitment and tissue damage during IAV infection.

### Immune cells have a glycocalyx that is shed following viral infection

Several recent studies have highlighted that, as well as endothelial cells, a range of immune cell populations have an HS glycocalyx that can regulate their function^15–20^. We therefore next analysed whether immune cells also shed their glycocalyx in response to IAV infection. We observed HS shedding from overall CD45^+^ cells (Fig. 4A), including monocytes (Fig. 4B) and neutrophils (Fig. 4C) 24 hours after IAV infection. To determine whether the immune cell glycocalyx was transcriptionally regulated across the broader IAV infection time course we re-analysed the single-cell IAV dataset from a glycocalyx remodelling and synthesis perspective. Consistent with our observed shedding, this revealed cell-specific and dynamic expression of *Hpse*, *Sulf1*, *Sulf2* and *Mmp2* (Fig. 4D). Interestingly monocytes and B cells were consistently the higher expressors of *Hpse*. We also observed immune cell-specific changes in GAG-related genes across the IAV infection time course (Fig. 4E), particularly in neutrophils and monocytes. There were also cell specific expression patterns across GAG and proteoglycan gene families (Fig. 4F), and in individual glycocalyx-associated genes (Supp. Fig. 5), consistent with time-dependent glycocalyx remodelling. We then applied flow cytometry-based glycocalyx analysis at 7 dpi (Fig. 4G). This showed recovery from early (24 hours post infection) HS shedding on tissue neutrophils by 7 dpi, sustained HS loss on monocytes, additional HS shedding from B cells, CD4^+^ and CD8^+^ T cells, and increased HS levels on alveolar macrophages. These distinct HS profiles suggest that immune cell glycocalyx remodelling is a regulated, cell-type-specific process rather than a passive consequence of inflammation.

**Figure 4.**
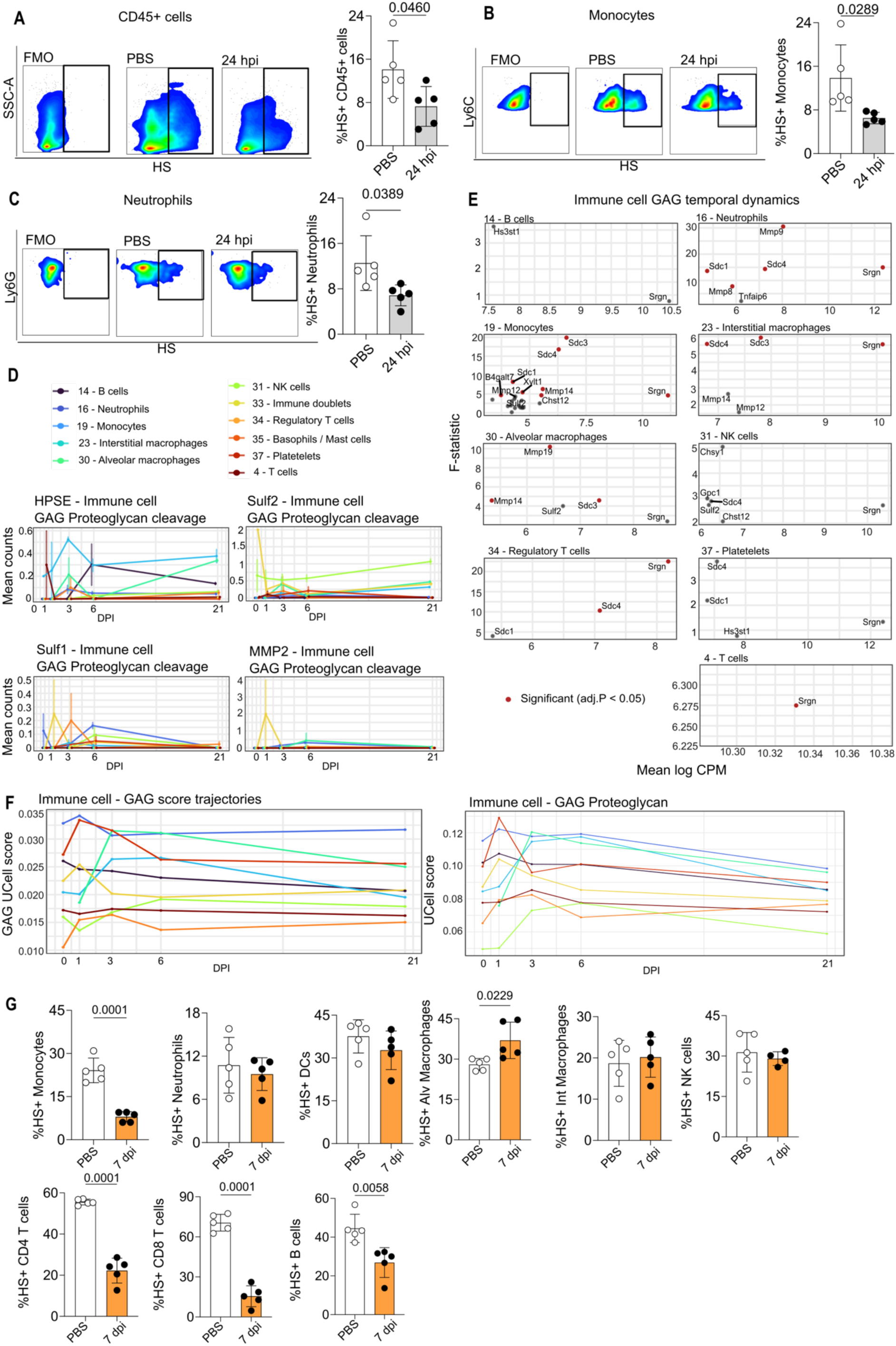
The immune cell glycocalyx is cell specifically re-modelled during the influenza immune response. Flow cytometry-based staining and quantification of HS on (A) overall immune cells (CD45^+^), (B) monocytes (Ly6C^hi^) and (C) neutrophils (Ly6G^+^) from the lungs of WT mice 24 hours after control or IAV infection. (D) Expression analysis of genes transcribing glycocalyx remodeling enzymes *Hpse*, *Sulf1*, *Sulf2* and *Mmp-2* in different immune cell subsets across the IAV infection time course. (E) Differential expression analysis of the most dynamic GAG synthesis genes across the IAV infection time course in immune cell subsets. (F) Expression analysis of “GAG glycan gene” and “proteoglycan protein core” grouped gene families. (G) Flow cytometry-based quantification of HS on different immune cells 7 days after control or IAV infection. Each dot represents an individual mouse, A-C and G (n=4-5), data are plotted as mean ± SD. Data from A-C and G were analysed by unpaired t-test with p values indicated.

**Figure 5.**
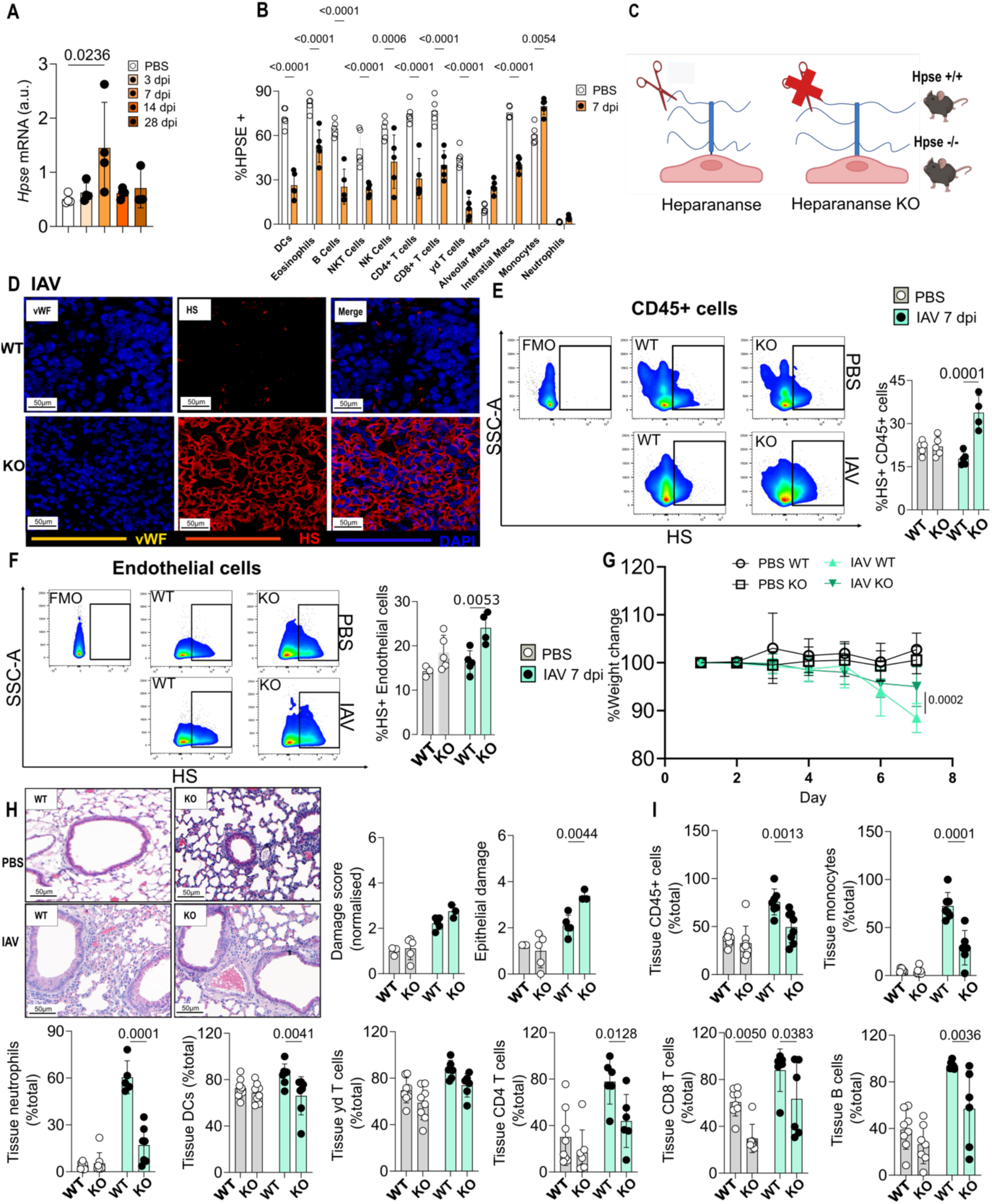
Heparanase drives immune cell recruitment during the viral immune response to influenza infection. (A) RT-qPCR analysis of whole lung expression of *Hpse* across the IAV infection time course. (B) Intracellular flow cytometry analysis of HPSE levels in different immune cells from the lungs 7 days post control (PBS) or IAV infection. (C) Schematic of HPSE mediated HS chain cleavage. (D) Representative FFPE Immunofluorescence staining of mouse lungs for cell nucleus (DAPI; blue), vWF (yellow) and HS (red) in immune cell rich areas. Scale bar is 50µm. Flow cytometry based staining and associated quantification of HS on (E) overall immune cells (CD45^+^) and (F) endothelial cells (CD45^-^CD31^+^); in lungs from WT and *Hpse* KO mice 7 days after control (PBS) or IAV infection. (G) Quantification of weight loss up to day 7 post IAV infection in WT and *Hpse* KO mice 7 days after control or IAV infection. (H) Haematoxylin and eosin (H&E) staining of mouse lung sections and associated overall histological tissue damage score and epithelial cell damage score. (I) Flow-cytometry based quantification of tissue (IV cd45^-^) immune cells (CD45+) and individual populations; in WT and *Hpse* KO mice 7 days after control or IAV infection. Each dot is an individual mouse, (A-B, E-F, H and J; n=3-5), (G and I, n=6-10), data are plotted as mean ± SD. Data from A were analysed by one-way ANOVA and Data from B-I were analysed by two-way ANOVA with p values indicated.

### Heparanase mediated glycocalyx shedding regulates immune cell recruitment and tissue damage following viral infection

Given our observation of endothelial, and immune, cell shedding of their glycocalyces following IAV infection we wished to determine the function of the enzyme heparanase (HPSE), which cleaves the HS glycan chain to mediate shedding^31^, in IAV infection. We first performed whole lung real time qPCR analysis, demonstrating increased expression of *Hpse* at day 7 dpi (Fig. 5A and Supp. Fig. 6A). We also undertook intracellular flow cytometry analysis of HPSE to reveal the presence of the protein in a range of immune cells at rest. Following IAV infection, HPSE^+^ populations were reduced in DCs, eosinophils, B cells, NK cells, NK T cells, CD4^+^ T cells, CD8^+^ T cells, TCRγδ cells, and interstitial macrophages at 7 dpi (Fig. 5B), presumably due to release into the vascular and tissue environment^16^. In contrast at this same time point there is an increase HPSE^+^ monocytes.

To mechanistically probe the function of this enzyme we used mice deficient in *Hpse* (Fig. 5C) and therefore unable to effectively shed the HS glycocalyx. We infected these mice with IAV and analysed their immune response, relative to WT controls, at 7 dpi. HS detection on tissue sections in non-vascular (VWF^-^) (Fig. 5D) and vascular (VWF^+^) (Supp. Fig. 6B) regions depicted a stark increase of HS in the *Hpse* KO murine IAV infected lungs. This immune cell (CD45^+^) (Fig. 5E), and endothelial cell (Fig. 5F), HS increase in the absence of HPSE was confirmed using flow cytometry. In contrast to the mice where HS accumulation capacity was removed (Fig. 3) the *Hpse* KO mice lost less weight (Fig. 5G) and limited overall tissue damage (Fig. 5H and Supp. Fig 6C), compared to WT IAV infected controls. Furthermore, the *Hpse* KO mice had significantly reduced levels of tissue CD45^+^ cells including tissue monocytes, neutrophils, DCs, CD4^+^ T cells, CD8^+^ T cells and B cells (Fig. 5I) in response to IAV infection, compared to WT controls at 7 dpi. Together confirming the importance of the glycocalyx in regulating immune cell recruitment during IAV infection.

Together the HPSE KO and eGC functional studies indicate that coordinated temporal remodelling of both vascular and immune cell glycocalyces regulates the balance between immune cell recruitment and tissue protection in murine IAV infection.

## Discussion

Here we reveal that IAV infection drives a coordinated, biphasic remodelling of the pulmonary glycocalyx that actively regulates the balance between immune cell recruitment and tissue damage. By developing a new tool to cell-specifically deplete sulfated GAGs (key glycocalyx components^10^), we demonstrate that remodelling is a previously unrecognised mechanism with broad implications for respiratory disease^2^.

Specifically, we observed that early (24 hours) after IAV infection, glycocalyx components are shed from both the endothelial-, and immune-cell surface. Using *Hpse* knock-out mice, that cannot cleave HS chains^39^, we demonstrated that this shedding was key to enable the entry of immune cells into the infected lungs from the circulation, a process vital to clearing infection^2^. Early eGC disruption during influenza infection is consistent with previous studies demonstrating that inflammatory mediators rapidly induce eGC degradation and shedding during acute inflammatory injury^10^. Whilst previous studies have demonstrated evidence of eGC shedding, detailed temporal assessment of glycocalyx remodelling within lung tissue has not been explored.

Subsequently, our longitudinal studies have revealed that the endothelial vasculature not only recovers the shed glycocalyx levels, compared to resting controls, but then accumulates glycocalyx components (HS, HA and SDC-1^8,9,14^) to levels higher than those seen at rest. Importantly, glycocalyx accumulation occurs alongside peak pulmonary inflammation and extensive immune cell infiltration. These findings were particularly striking as they contrast with the commonly proposed model of progressive glycocalyx loss during inflammatory disease^8–10,24^. Instead, these data suggest that IAV infection induces active eGC remodelling rather than uniform glycocalyx destruction alone.

Our new mouse model, to cell specifically deplete glycocalyx components (GAGs), revealed that accumulation of the vascular glycocalyx is key in defining the tissue impact of the immune response. Endothelial specific removal of the key GAG synthesis gene *B4Galt7*^29^ resulted in excessive immune cell recruitment and tissue damage. As a result, our data demonstrate that accumulation of the eGC barrier at later time points of murine IAV infection limits excessive immune cell recruitment to control tissue damage and enable restoration of tissue homeostasis.

We also observed distinct temporal regulation across individual glycocalyx-associated components on endothelial cells. Whilst HS and HA on the endothelial surface demonstrated early reduction at 24 hours post-infection followed by later accumulation during peak disease, SDC-1 displayed a different biphasic pattern characterised by reduced expression at 3 dpi followed by increased expression at 7 and 14 dpi. These findings suggest that proteoglycan core proteins and associated GAG chains undergo distinct temporal regulation during influenza-induced pulmonary inflammation. This may be due to the early shedding or degradation of HS GAG chains exposing the SDC-1 core protein to subsequent proteolytic cleavage and shedding from the endothelial cell surface, consistent with previous studies demonstrating sequential glycocalyx degradation during inflammatory injury^40^. Furthermore, across the IAV infection time-course (7-14 dpi) the glycocalyx is dynamically and cell specifically remodelled across diverse immune cell populations, where glycocalyx analysis has previously been limited^9,16–21^. Together, these data demonstrate that the glycocalyx is temporally regulated, playing cell-specific functions throughout the viral immune response.

This study updates the existing paradigm that inflammation simply induces glycocalyx shedding to reflect endothelial injury^8–10,24,25^. We now propose a model in which the pulmonary eGC is first shed before re-forming to a greater extent rather than simply returning to its homeostatic form. As such the endothelial, and immune, cell glycocalyces are active participants in the host defence against IAV infection and in the subsequent tissue repair process. Given the importance of glycocalyx biology across vascular beds and tissue contexts, these findings may have broad implications for understanding immune regulation in respiratory and wider inflammatory diseases.

## Acknowledgements

We thank Dr David Boyd for generously sharing the murine influenza infection longitudinal single cell sequencing dataset that was re-analysed in this study. We acknowledge the help and input from the genome editing unit and bioimaging core facility at the University of Manchester.

## Authors Contributions

Conceptualization: IZN, DPD

Methodology: IZN, IA, RD, JP, AJ, AJLR, ZZ, ADA, KOL

Investigation: IZN, IA, RD, JP, AJ, AJLR, ZZ, ADA, KOL

Visualization: IZN, DPD

Resources: JEA, JPL

Funding acquisition: JEA, TH, DPD

Project administration: DPD

Supervision: TH, DPD

Writing – original draft: IZN, DPD

Writing – review & editing: All

## Data, code, and materials availability

Any data not explicitly evident in the paper will be available on reasonable request. Single-cell data are available on the NCBI Short Read Archive under BioProjects PRJNA612345 and PRJNA613670.

## List of Supplementary Materials

Materials and Methods

Tables S1-5

Figs. S1 to S6

