## Supplementary information for "Biphasic glycocalyx remodelling balances immune cell recruitment and tissue damage during influenza infection"

#### Materials and Methods

##### Mice

All animal experiments were locally and ethically approved and performed in accordance with the UK Home Office Animals (Scientific Procedures) Act of 1986 under project licenses PP0286419 or P5636795 granted by the UK Home Office and approved by the University of Manchester Animal Welfare and Ethical Review Body. Euthanasia was performed by intraperitoneal pentobarbitone overdose. Female and male mice aged 8-12 weeks were used for all experiments. Animals were housed under specific pathogen-free (SPF) conditions in individually ventilated cages with a maximum of five mice per cage and maintained under a 12-hour light/dark cycle at controlled temperature (20-24°C) and humidity (45-65%). Standard rodent chow and water were provided ad libitum, together with environmental enrichment consisting of nesting material, cardboard shelters and gnawing blocks. Animals were acclimatised for a minimum of one week before experimental procedures commenced. All genetically modified mice were bred and maintained within the University of Manchester Biological Services Facility.

Global heparanase knockout (Hpse−/−) mice on a C57BL/6J background were used to investigate the role of heparan sulphate degradation during influenza A virus infection. Hpse−/− mice were originally generated through targeted disruption of the Hpse gene as previously described[1]. Wild-type (WT) littermates were used as experimental controls. *B4galt7* fl/fl mice were generated as described below and crossed with tamoxifen-inducible *Cdh5*-CreERT2 mice to generate endothelial cell-specific *B4galt7* conditional knockout mice. Cre-negative *B4galt7* fl/fl littermates were used as controls. Breeding colonies were maintained by heterozygous matings and offspring were genotyped prior to experimental use. Genomic DNA was isolated from ear punch or tail biopsy tissue, and genotyping was performed by Transnetyx. Female and male C57BL/6J mice aged 8–12 weeks were used for all WT IAV time-course experiments unless otherwise stated. Animals were obtained from Charles River.

**Influenza A virus (IAV) infection model**

Influenza A virus (IAV) infection was performed using the mouse-adapted A/Puerto Rico/8/1934 (H1N1) (PR8) strain. Mice were anaesthetised using 4% isoflurane for induction and maintained under 2% isoflurane with oxygen at 1 L/min during virus administration. Anaesthetised mice were infected intranasally with 18 plaque-forming units (PFU) of PR8 influenza virus in a total volume of 25 µL PBS, while control mice received 25 µL PBS by the same route. Mice were weighed daily for welfare monitoring and assessment of disease progression. Percentage body weight was calculated relative to each animal's pre-infection weight. Clinical observations included assessment of general appearance, posture, activity, grooming behaviour and respiratory signs.

**Generation of a conditional *B4galt7* KO allele**

A conditional KO allele of the mouse *B4galt7* gene was generated using CRISPR-Cas9 mediated homology-directed repair (HDR) to introduce loxP sites flanking the critical exon 3. Guide RNAs (sgRNAs) were designed to target predicted non-conserved intronic regions flanking exon 3, selected to minimise predicted off-target activity and to ensure disruption of the sgRNA target sites following successful HDR. Two sgRNAs were used: g525 (5′-CAGGCTGTCAAGTTCCACCT-3′) and g684 (5′-ACAGCCTGTAAAGCGGGGCC-3′), synthesised as chemically modified sgRNAs (IDT).

*Long single-stranded DNA donor template design and production*

HDR was mediated using a long single-stranded DNA (lssDNA) donor template containing loxP sites positioned 5′ and 3′ of exon 3. The donor also incorporated an *EcoRI* restriction site adjacent to the 5′ loxP and a *BamHI* site adjacent to the 3′ loxP to facilitate molecular genotyping. Homology arms of approximately 85 bp were used on each side of the insert.

The lssDNA donor was generated using a previously described in-house method[2]. Briefly, a biotinylated forward primer and a gene-specific reverse primer were used to PCR-amplify a biotin-labelled double-stranded DNA product. The PCR product was immobilised on a streptavidin column and denatured using NaOH to remove the non-biotinylated strand. A hybridisation primer was then annealed to generate a short double-stranded region, allowing cleavage with *SphI* to release purified, high-quality lssDNA. The final donor DNA was concentrated and Sanger sequenced prior to embryo microinjection.

sgRNAs were resuspended in RNase-free injection buffer (1 mM Tris-HCl, pH 7.5; 0.1 mM EDTA) to a concentration of 1 µg/µl. For ribonucleoprotein (RNP) complex formation, sgRNAs were mixed with recombinant Engen Cas9 protein (NEB) and incubated at room temperature for 10 min. The final injection mix contained sgRNA–Cas9 RNPs (40 ng/µl each), Cas9 mRNA (20 ng/µl), and lssDNA donor template (20 ng/µl).

Cryopreserved one-cell C57BL/6J embryos generated by *in vitro* fertilisation were thawed and cultured prior to microinjection. The CRISPR-Cas9 injection mix was delivered by pronuclear microinjection into zygotes over two consecutive days. Injected embryos were cultured overnight to the two-cell stage and surgically transferred into the oviducts of day 0.5 post-coitum pseudopregnant recipient females. Resulting pups (15 total) were allowed to develop to weaning age before genotyping.

For genotyping, genomic DNA was extracted from ear biopsies using the REDExtract-N-Amp Tissue PCR Kit (Sigma-Aldrich). PCR was performed using primer pairs flanking the 5′ (ctF1/ctR1) and 3′ (ctF2/ctR2) loxP integration sites, with all primers designed for a melting temperature of approximately 60 °C. PCR products were analysed by Qiaxcel capillary electrophoresis (Qiagen).

Successful loxP integration candidates were initially identified by an increase of 40 bp in amplicon size and by diagnostic restriction enzyme digestion. PCR products were digested directly without purification using *EcoRI* (5′ loxP) or *BamHI* (3′ loxP) in CutSmart buffer at 37 °C for 1 h and analysed by electrophoresis to confirm likely loxP insertion. This analysis indicated 3 pups to have integrated the 5’ loxP only and one pup to likely have both 5’ and 3’ loxP integrated, which was taken forward for further analysis.

The putative founder was further validated by high-fidelity PCR amplification (KOD polymerase: Sigma) using primers DD03_ctF1 and DD03_ctR2, cloning into pCR-Blunt vectors, and Sanger sequencing. Sequencing confirmed the presence of both loxP sites in cis on a single allele, generating a fully floxed *B4galt7* allele. A 128 bp intronic deletion was identified between the loxP sites (Fig. S4E), consistent with non-homologous end joining following Cas9 cleavage; this deletion is not predicted to disrupt gene function.

The confirmed founder was bred to wild-type C57BL/6J mice to assess germline transmission. F1 offspring were genotyped using the same PCR strategy, and correct transmission of the loxP flanked allele was confirmed by PCR and sequencing. The recommended nomenclature for the allele is **C57BL/6J.*B4galt7*** $^{\mathbf{Em1Uman}}$.

**Supplementary Table 1. Primers used for genotyping**

| Primer name | Primer sequence |
|---|---|
| DD03_ctF1 | ACCATCACCTCTTGCCTCTC |
| DD03 ctR1 | ACCACCCAGCACTAATGTCA |
| DD03_ctF2 | CTAGGCAGTTGGGAGGGTAG |
| DD03_ctR2 | GTCCCATCAAGAGCCAGTGT |

**Final allele sequence**

Capital letters = LoxP

Purple = exon 3

Strikethrough = deleted 128nt

Remaining sequence is intronic

ggagcccgatctcccagcttccataggatcactcactcaggctgtcaagttccaATAACTTCGTATAGCAT
ACATTATACGAAGTTATgaattcccttggatgttgcctgccattctggccctcctggcagccctgtctaggaga
atagctgcccacctgccctgccagtttccatttcaatacctggcttgacttgctagagaacatggatttgtttctcgctctttct
tccatgtattttgaacagtttctggcataaaccttcgtgcagcaagcagagggggcggtgttccacagtgacattagtgc
tgggtggtggcagcaaagagactcatgcatagccagagtctgagctaggttgataaacaagcttgcttcctctcccta
gcatcgtttatcttgtataggttcaatcgggcagcactcatcaatgtgggcttcctggagagcagcaacagcacagact
acatcgccatgcacgatgtggacctgctccctctcaatgaggagctggactatggcttcccggaggctgggcccttcc
atgtggcctccccagagctccaccctctctaccactacaagacctatgtgggcggcattctgctgctgtccaaacagca
ctaccagctggtgaggcccagccccacatgctctgctcatttgctggggatcttccccatcgtgggtctaggcagttggg
agggtagagcagccttcctggggcccagcagatggagcagtgccagagcctgagcagcagccagagactgtgag
ggaaccaggagggagcacttagccagggagagctctgctgggaccacagaagccccagtgtcagctggtcATA
ACTTCGTATAGCATACATTATACGAAGTTAT~~ggatcccagggacctagaggtgggaaggcctgc~~
~~agaacctccccctccctgcaagggtaccaagtcttcctttctcttcccaggcctcagcgcctgcccctcctgtgagaag~~
~~gctgtgtagctacagcctgtaa~~agcggggcctggttggggagtgggggatggagctcctccctcccc

**Tamoxifen-induced endothelial-specific B4galt7 knockout mice**

Mice carrying loxP-flanked (*B4galt7*fl/fl) alleles were crossed with tamoxifen-inducible *Cdh5*-CreERT2 transgenic mice (EMMA) to generate endothelial-specific *B4galt7*fl/fl *Cdh5*-CreERT2 KO animals. Littermates lacking the Cre transgene (*B4galt7*$^{fl/fl}$) were used as WT controls throughout the study.
To induce endothelial-specific recombination, mice received tamoxifen (Sigma) dissolved in 100% ethanol and corn oil (Sigma) at a dose of 2 mg per mouse by intraperitoneal injection once daily for five consecutive days. Following the final tamoxifen administration, mice were left for 7 days to allow time for Cre-mediated recombination before influenza A virus infection. Successful recombination led to deletion of *B4galt7* specifically within vascular endothelial cells while preserving gene expression in non-endothelial cell populations. Successful endothelial-specific deletion of *B4galt7* was confirmed by PCR, flow cytometry and immunofluorescence.

**Tissue collection**

At the designated experimental endpoint, mice were euthanised with 3 mg intraperitoneal pentobarbitone then exsanguinated via the femoral artery and blood was collected. Blood samples were allowed to clot at room temperature before centrifugation at 300 x g for 5 min, and the resulting serum was collected and stored at -80°C until further analysis. Following blood collection, the thoracic cavity was opened to expose the lungs and heart. Whole lungs were inflated with 2 mL inflation enzyme solution (1 mg/mL Dispase II (Roche UK), and 50 µg/mL DNase I, (Roche, UK)) via an intratracheal cannula. The lungs were carefully excised and processed according to the downstream application. For flow cytometry, lung tissue was immediately transferred into ice-cold inflation enzyme solution. For RNA analysis, lung tissue was preserved in RNAlater (Invitrogen) before storage at -80°C. For histological analyses, lungs were inflation-fixed with 10% neutral buffered formalin (NBF) at a constant pressure before immersion fixation. Following fixation, tissues were processed for paraffin embedding prior to sectioning and staining.

**Lung tissue digestion and single-cell suspension preparation**

Single-cell suspensions were prepared from whole lung tissue for subsequent flow-cytometric analysis of immune and endothelial cell populations. Immediately following tissue collection, whole lungs were transferred to the same inflation enzyme solution used during tissue collection and incubated at 37°C for 1 h. Lung lobes were subsequently separated and incubated at 37°C for 10 min with 0.1 mg/mL Dispase II (Roche), Liberase TL (Roche) and 50 µg/mL DNase I (Roche), after which enzymatic digestion was terminated using 5 mM EDTA. Following digestion, the tissue was mechanically dissociated by gentle trituration using a pipette and passed through a 70 µm cell strainer to generate a single-cell suspension.

Cells were collected by centrifugation at 600 × g for 6 min at 4°C. Red blood cells were removed using ACK (ammonium-chloride-potassium) Red Blood Cell Lysis Buffer (Gibco) according to the manufacturer's instructions. Cells were washed with PBS and resuspended in FACS buffer consisting of PBS supplemented with 2% FBS and 2 mM EDTA. Viable cells were counted manually using a haemocytometer with trypan blue exclusion where required.

**Flow cytometry**

Approximately 1-5 × $10^6$ cells were transferred into individual wells of a 96-well V-bottom plate for antibody staining. Unless otherwise stated, washing steps were performed by adding 200 µL FACS buffer followed by centrifugation at 600 × g for 3 min at 4°C. Cell viability was assessed using Zombie UV viability dye (BioLegend) according to the manufacturer's instructions. Fc receptors were blocked using purified anti-mouse CD16/CD32 (BD Biosciences) for 15 min at 4°C to minimise non-specific antibody binding. Cells were subsequently incubated with fluorophore-conjugated antibodies against the relevant surface markers (Supplementary Table 2) for 30 min at 4°C in the dark. For intracellular detection of heparanase (HPSE), surface staining was completed before fixation and permeabilisation using the FOXP3/Transcription Factor Staining Buffer Set (eBioscience) according to the manufacturer's instructions. Cells were incubated overnight at 4°C with anti-heparanase primary antibody diluted in permeabilisation buffer. Following primary-antibody incubation, cells were washed twice in permeabilisation buffer and incubated with the appropriate fluorophore-conjugated anti-rabbit IgG secondary antibody. Cells were washed before final resuspension in FACS buffer for acquisition. Data were acquired on a BD Fortessa flow cytometer (BD Biosciences) using FACSDiva software and analysed using FlowJo (Tree Star, USA). Samples were acquired to a target of approximately 1 × $10^6$ events per sample where cell yield permitted. Compensation was performed using single-stained compensation controls prepared using UltraComp eBeads (Thermo Fisher Scientific) for each fluorochrome included within the experimental panel. Fluorescence-minus-one (FMO) controls were included where required to assist with gate placement. Representative gating strategies for endothelial and immune-cell populations are shown in Supplementary Fig. S1A-B.

For absolute cell quantification, Thermo Fisher Scientific counting beads were used. For each sample, 25 µL of counting beads was diluted in 175 µL FACS buffer and combined with 200 µL of cell suspension prior to acquisition. The bead preparation contained 0.52 × $10^6$ beads per 50 µL; therefore, 25 µL corresponded to 2.6 × $10^5$ beads added per sample. Absolute cell numbers were calculated from the ratio of gated cellular events to acquired bead events using:

absolute cell number = (cell events / bead events) × 2.6 × $10^5$

Absolute cell numbers were subsequently normalised to the recorded lung weight and expressed as cells per gram of lung tissue.

For detection of cell-surface heparan sulphate (HS), anti-heparan sulphate monoclonal antibody 10E4 was used in an optimised staining protocol. Cells were incubated with biotinylated 10E4 overnight, washed as described above and subsequently incubated with fluorophore-conjugated streptavidin for 1 h. For detection of cell-surface hyaluronan (HA), cells were incubated with hyaluronan-binding protein (HBP; Merck; 1:200), washed and subsequently incubated with fluorophore-conjugated streptavidin for 1 h before flow-cytometric acquisition. Reagent dilutions are provided in Supplementary Table 2.

To distinguish vascular-accessible leukocytes from leukocytes localised within the lung tissue, intravascular CD45 labelling was performed immediately prior to euthanasia. Mice received fluorophore-conjugated anti-mouse CD45 antibody (clone 30-F11; APC; BioLegend) by intravenous tail-vein injection 3 min before tissue collection. Cells positive for intravenously administered CD45 (ivCD45+) were classified as vascular-accessible leukocytes, whereas cells negative for intravenous CD45 but positive for CD45 following ex vivo staining (ivCD45−CD45+) were classified as tissue-localised leukocytes.

**Supplementary Table 2. Flow cytometry antibodies and detection reagents used to characterise murine samples**

| Antibody | Clone | Dilution | Conjugate | Company |
|---|---|---|---|---|
| CD45 | 30-F11 | 200 | BV785 | BioLegend |
| CD31 | 390 | 200 | BV711 | BD Optibuild |
| EpCAM | G8.8 | 200 | PE | BioLegend |
| SDC-1 (CD138) | 28-1-2 | 200 | PE/Cy7 | BioLegend |
| WGA | - | 2000 | FITC | |
| HS | 10E4 | 200 | SA | AMSBIO |
| HBP | - | 200 | SA | Merck life science |
| Podoplanin | 8.1.1 | 200 | PerCP/Cy5.5 | BioLegend |
| AckR1 | - | 200 | FITC | - |

| | | | | |
|---|---|---|---|---|
| CD4 | RM4-5 | 200 | FITC | BioLegend |
| CD8 | 53-6.7 | 800 | PerCP/Cy5.5 | BioLegend |
| MerTK | 2BIOC42 | 200 | APC | BioLegend |
| Ly6C | HK1.4 | 200 | AF700 | BioLegend |
| Ter119 | Ter119 | 200 | APC 780 | eBiosciences |
| CD3 | 17A2 | 100 | APC 780 | eBiosciences |
| CD11c | N418 | 800 | BV605 | BioLegend |
| TCRb | H5-597 | 200 | APC 780 | eBiosciences |
| Ly6G | 1A8 | 100 | BV510 | BioLegend |
| B220 | RA3-6B2 | 100 | BV650 | BioLegend |
| CD11b | M1/70 | 800 | BV711 | BioLegend |
| CD64 | X54-5/7.1 | 100 | PE | BioLegend |
| Siglec F | CF-e50-2440 | 400 | PECF594 | BioLegend |
| NK1.1 | PK136 | 200 | PE/Cy5 | eBiosciences |
| TCRγδ | eBIOGL3 | 200 | PE/Cy7 | eBiosciences |
| Zombie UV | - | 2000 | UV | BioLegend |
| SA | - | 500 | PECF594 | BioLegend |
| ivCD45 | 30-F11 | 3 μg | APC | BioLegend |
| HPSE | 24529-1-AP | 200 | - | Proteintech |
| CD19 | (1D3) 6D5 | 200 | APC/eFluor780 | BioLegend |
| MHC II | M5/114.15.2 | 20000 | PE-Cy5 | BioLegend |

*Cell Sorting*

For validation of endothelial-specific recombination, lung single-cell suspensions were prepared as described above from Cre-positive tamoxifen-treated, Cre-positive vehicle-treated, Cre-negative tamoxifen-treated and Cre-negative vehicle-treated mice. Cells were stained with viability dye and antibodies against CD45, CD31 and EpCAM. Live singlets were identified before isolation of endothelial cells as CD45−CD31+EpCAM− and immune cells as CD45+. Cell populations were sorted using a BD FACS Aria cell sorter (BD Biosciences) and subsequently processed for genomic DNA extraction and PCR analysis of B4galt7 recombination. Representative sorting strategies are shown in Supplementary Fig. S4B-C.

#### Histology and immunofluorescence of lung

*Tissue preparation*

Lung tissue collected for histological analysis was processed as formalin-fixed, paraffin-embedded (FFPE) tissue. Following euthanasia, the trachea was cannulated and lungs were inflated intratracheally with 1 mL of 10% neutral buffered formalin (NBF; Sigma) at constant pressure to preserve pulmonary architecture. The trachea was secured to maintain inflation and the intact lungs were excised and immersed in excess 10% NBF for 24 h at room temperature. Tissues were subsequently transferred to 70% ethanol for 24 h before routine dehydration and processing using an ASP300S automated tissue processor (Leica). Lungs were orientated consistently during paraffin embedding to facilitate sectioning of comparable anatomical regions between animals. FFPE blocks were sectioned at 5 µm using a rotary microtome (Leica Biosystems), floated on a 40–45°C water bath and mounted onto Superfrost Plus microscope slides (Epredia). Slides were dried overnight at 37°C prior to staining.

*H&E staining*

Haematoxylin and eosin (H&E) staining was performed to assess pulmonary tissue architecture and influenza-induced lung pathology. FFPE lung sections (5 µm) were deparaffinised, rehydrated through graded alcohols and stained using an automated Shandon Varistain™ 24-4 slide stainer (Thermo Shandon Ltd., UK) according to the manufacturer's standard staining programme. Following staining, sections were dehydrated through ascending concentrations of ethanol, cleared in xylene and coverslipped. Whole-slide brightfield images were acquired using a 3D HISTECH Pannoramic-250 slide scanner under identical imaging settings for all experimental groups.

*Immunofluorescence staining*

Immunofluorescence staining was performed on 5 µm FFPE lung sections. Sections were deparaffinised in xylene and rehydrated through decreasing concentrations of ethanol into distilled water. Heat-induced antigen retrieval was performed using Tris-EDTA buffer (10 mM Tris, 1 mM EDTA, 0.05% Tween-20, pH 9.0) for 20 min. Sections were subsequently permeabilised using 0.5% Triton X-100 (Sigma-Aldrich) in Tris-buffered saline (TBS; 50 mM Tris, 150 mM NaCl, pH 7.6).

Non-specific antibody binding was blocked using donkey serum diluted in 1% BSA in TBS for 30 min at room temperature. Endogenous biotin activity was blocked using an Avidin/Biotin Blocking Kit (Vector Laboratories) according to the manufacturer's instructions. As mouse primary antibodies were used on mouse tissue, additional blocking was performed using Mouse Ig Blocking Reagent from the Mouse on Mouse (M.O.M.) Immunodetection Kit (Vector Laboratories) for 1 h at room temperature.

Primary antibodies (Supplementary Table 3) were diluted in M.O.M. diluent and incubated with tissue sections overnight at 4°C in a humidified chamber. Where mouse-on-mouse primary antibodies were used, including anti-HS clone 10E4, sections were washed in TBS and incubated with the biotinylated anti-mouse secondary antibody supplied with the M.O.M. Immunodetection Kit for 10 min. HS was subsequently detected using fluorophore-conjugated streptavidin for 1 h at room temperature. Rabbit primary antibodies, including vWF and NP, were detected using the appropriate fluorophore-conjugated donkey anti-rabbit secondary antibody.

Nuclei were counterstained with DAPI (4′,6-diamidino-2-phenylindole; Thermo Fisher Scientific) at a final concentration of 0.2 μg/mL for 5 min. Following final washes, sections were mounted using ProLong Gold Antifade Mountant (Thermo Fisher Scientific) and coverslipped.

**Whole-organ lung clearing and light-sheet microscopy**

For whole-organ visualisation of the pulmonary vascular glycocalyx, mice received intravenous administration of directly conjugated antibodies against CD31 and heparan sulphate (HS) prior to tissue collection. Anti-CD31 antibody (clone 390) conjugated to FITC was administered at 15 μg per mouse, together with 4 μg of Alexa Fluor 555-conjugated anti-HS antibody (clone F58-10E4). Antibodies were allowed to circulate for 30 min before animals were perfused and lungs were fixed with 4% paraformaldehyde (PFA).

Fixed lungs were subsequently cleared using the F-DISCO tissue-clearing method and prepared for whole-organ light-sheet fluorescence microscopy. As both antibodies were directly fluorophore-conjugated, no secondary-antibody staining was required. Cleared lungs were imaged using a Zeiss Lightsheet Z.1 and image data were visualised and reconstructed using Imaris software.

**Supplementary Table 3. Antibodies and detection reagents used for immunofluorescence**

| Antibody | Clone | Dilution | Company |
|---|---|---|---|
| vWF | Rb-x | 1:100 | Merck |
| HS | F58-10E4 | 1:250 | AMSBIO |
| HBP | - | 1:200 | Merck |
| SA 637 | - | 1:800 | BD Biosciences |
| SA 557 | - | 1:800 | BD Biosciences |
| Donkey-anti-rabbit 637 | polyclonal | 1:200 | Bio-Techne |
| Donkey-anti-rabbit 557 | polyclonal | 1:200 | Bio-Techne |

**Immunofluorescence image Analysis**

Whole-slide fluorescence images were acquired using a 3D HISTECH Pannoramic-250 slide scanner at ×20 magnification. Whole-slide images were imported into QuPath version 0.5.1, where individual fluorescence channels were visualised and identical display settings were applied across all experimental groups within each experiment. Representative images were selected to illustrate the spatial localisation of HS, HA and vWF within the pulmonary vasculature.

For selected localisation analyses, fluorescence-intensity profiles were generated across defined regions of interest using the RGB Profiler plugin in ImageJ version 1.53a to assess the relative spatial localisation of glycocalyx-associated and endothelial markers.

Influenza nucleoprotein (NP) staining was analysed using QuPath version 0.5.1. Whole-slide images were imported into QuPath and the whole left lung lobe was manually annotated prior to analysis. A positive-pixel classifier was applied using identical threshold settings across all sections within an experiment to identify NP-positive staining. Pulmonary viral antigen burden was quantified as the percentage of NP-positive tissue area relative to the total annotated lung area. The same classifier parameters were applied to all slides within each experiment to ensure consistency between experimental groups. Image files were coded and analysis was performed blinded to genotype and treatment group until quantitative analysis had been completed.

**Histopathological scoring**

Whole-lung sections were examined using QuPath version 0.5.1, and pathological changes were assessed across the entire tissue section to ensure representative evaluation of each specimen. Pulmonary pathology was assessed using a semi-quantitative scoring system adapted from previously published influenza histopathology approaches[3,4]:

Six characteristic features of influenza-induced lung injury were assessed: inflammatory-cell infiltration, peribronchial inflammatory-cell infiltration, epithelial-cell damage, alveolar septal thickening, pulmonary oedema and pulmonary haemorrhage. Each parameter was assigned a severity score from 0 to 4, where 0 indicated no pathological change, 1 mild involvement, 2 moderate involvement, 3 severe involvement and 4 very severe involvement. Individual parameter scores were summed to generate an overall histopathology score for each animal, with higher scores indicating greater lung injury. Slides were coded prior to assessment and histopathological scoring was performed blinded to experimental group and genotype by three independent researchers. The mean score from the three assessors was used for subsequent analysis.

**Single cell transcriptomics analysis**

*Dataset and experimental design*

Single-cell RNA sequencing data were analysed from a pre-processed, SCTransform-normalised Seurat object containing cells from a mouse influenza A/PR8 infection time-course[5,6]. The dataset followed a cross-sectional design, with five animals pooled at each of five timepoints: 0, 1, 3, 6 and 21 days post-infection. Cells from all timepoints were aggregated into a single Seurat object for downstream analysis. Mouse of origin was resolved from hashtag oligo (HTO) labelling using Seurat's demultiplexing algorithm, and only HTO singlets were retained for animal-level pseudo-bulk analyses used for statistical validation.

Cluster identities were inherited from prior processing using Louvain clustering, resulting in 38 clusters. No additional re-clustering was performed. Clusters were annotated using established marker genes and subsequently grouped into four broad lineage categories: Immune, Epithelial, Mesenchymal and Endothelial. These broad cell-type classifications were used throughout the analysis to support lineage-level comparisons.

Four clusters were excluded from downstream quantitative analyses due to quality concerns but were retained in overview UMAP visualisations for reference: cluster 9, which showed high mitochondrial content; cluster 33, annotated as immune doublets; cluster 36, annotated as multiplets; and cluster 37, annotated as platelets.

*Gene set curation and single-cell scoring*

Glycocalyx-related gene categories, including proteoglycans, glycosaminoglycan (GAG) biosynthesis enzymes, degradation enzymes and glycoside hydrolases, were curated from a previous publication[7]. Gene-set activity was quantified at single-cell resolution using UCell, a rank-based gene signature scoring method.

A composite glycocalyx score was generated from the union of all glycocalyx-associated genes. Additional composite biosynthesis and degradation scores were generated from genes belonging to the corresponding functional categories (Supplementary Table 4).

Initial glycocalyx analyses revealed marked heterogeneity across broad cell types, with particularly prominent signatures within Immune and Endothelial populations. To examine these lineage-specific patterns in greater detail, a focused analysis was performed using eight curated GAG-related gene sets comprising proteoglycan core proteins, proteoglycan-cleaving enzymes, overall GAG biosynthesis enzymes, heparan sulphate-specific biosynthesis enzymes, chondroitin sulphate-specific biosynthesis enzymes, hyaluronan synthesis enzymes, and heparan sulphate and chondroitin sulphate sulphation enzymes. These gene sets were designed ourselves and scored using UCell. An overall GAG score was calculated from the union of genes across all eight categories.

*Visualisation of glycocalyx and GAG programmes*

Gene-set activity was visualised using UMAP feature plots, boxplots of UCell scores and hierarchically clustered heatmaps of per-cluster mean UCell scores.

Heatmaps were displayed using two complementary scaling approaches. Column-wise z-scoring was used to identify clusters with relatively high or low activity for a given programme, whereas row-wise z-scoring highlighted programmes that were relatively enriched within individual clusters.

Expression of representative marker genes was visualised using dot plots, where dot size represented the proportion of cells expressing a gene and dot colour represented mean scaled expression.

*Correlation analysis*

Relationships between glycocalyx- and GAG-associated transcriptional programmes were examined by calculating correlations between UCell scores. These analyses were used to identify coordinated transcriptional signatures across cells, clusters and broad cell types and to place lineage-specific glycocalyx patterns within a broader biological context.

*Temporal dynamics: pseudobulk spline modelling*

To investigate temporal changes in glycocalyx-associated pathways during infection, a pseudobulk spline-modelling approach was applied to genes contained within the curated GAG gene sets.

Treating individual cells as independent replicates would inflate the statistical significance of any temporal trend, so cells classified by HTO demultiplexing as doublets or negative were excluded, and raw RNA counts from the remaining HTO singlets were summed per animal at each timepoint for each QC-passing cluster. This generated one pseudobulk expression profile per animal per timepoint, giving genuine biological replicates and statistically valid p-values for the temporal model.

Clusters were excluded if they contained fewer than three timepoints or fewer than four independent pseudobulk replicates. Normalisation and gene-expression filtering were applied across the full transcriptome rather than the GAG panel alone, so that gene-level statistics were calibrated against genome-wide variation before GAG-specific results were extracted for reporting.

Gene expression was modelled as a function of days post-infection using natural cubic splines within the limma-voom framework, with spline complexity adjusted according to the number of available timepoints for each cluster.

Genes were ranked within each cluster using a moderated F-statistic, reflecting the strength of temporal variation across the infection time-course and flagged as statistically significant at a Benjamini-Hochberg adjusted p-value below 0.05. The top 10 genes per cluster were retained, and genes appearing among the top-ranked genes of multiple clusters were highlighted as candidates displaying recurrent temporal behaviour across distinct cellular contexts.

The temporal modelling was used to identify genes displaying dynamic expression patterns across infection and to provide temporal context for glycocalyx- and GAG-associated transcriptional programmes identified by UCell scoring.

*Gene expression trajectories*

Temporal expression trajectories were generated for genes selected from the spline analysis. Gene selection was performed across all glycocalyx and GAG gene sets, retaining genes that appeared among the top-ranked temporal candidates in at least one cluster.

Mean expression and standard error of the mean (SEM) were calculated per cluster at each timepoint from per-animal mean expression values derived from RNA-assay count data (HTO singlets only), so that SEM reflected variation across biological replicates rather than across individual cells. Trajectories were also displayed separately for Immune, Endothelial, Epithelial and Mesenchymal populations, with individual clusters shown as separate curves within each broad lineage category.

One plot was generated for each gene and broad cell-type combination. These plots were used to examine temporal expression patterns underlying lineage-specific glycocalyx and GAG transcriptional programmes and to aid interpretation of the broader UCell-based findings.

*Software*

Analyses were performed in R (v4.4.3) using Seurat (v5.4.0)[8], UCell (v2.10.1)[9], limma (v3.62.2)[10], edgeR (v4.4.2)[11] and ComplexHeatmap (v2.22.0)[12]. Data manipulation and visualisation were performed using packages from the tidyverse ecosystem together with ggplot2, patchwork, ggrepel and ggridges[13].

**Supplementary Table 4. Gene family information.**

| Family | Gene |
|---|---|
| Proteoglycan | *Sdc1* |
| | *Sdc2* |
| | *Sdc3* |
| | *Sdc4* |
| | *Gpc1* |
| | *Gpc2* |
| | *Gpc3* |
| | *Gpc4* |
| | *Gpc5* |
| | *Gpc6* |
| | *Hspg2* |

| | |
|---|---|
| | *Srgn* |
| Proteoglycan cleavage | *Hpse*1 |
| | *Hpse*2 |
| | *Mmp1* |
| | *Mmp2* |
| | *Mmp3* |
| | *Mmp4* |
| | *Mmp5* |
| | *Mmp6* |
| | *Mmp7* |
| | *Mmp8* |
| | *Mmp9* |
| | *Mmp10* |
| | *Mmp11* |
| | *Mmp12* |
| | *Mmp13* |
| | *Mmp14* |
| | *Mmp15* |
| | *Mmp16* |
| | *Mmp17* |
| | *Mmp18* |
| | *Mmp19* |
| | *Mmp20* |
| | *Mmp21* |
| | *Mmp22* |
| | *Mmp23* |
| | *Mmp24* |
| | *Mmp25* |
| | *Mmp26* |
| | *Mmp27* |
| | *Mmp28* |
| | *Tnfaip6* |
| | *Sulf1* |
| | *Sulf2* |
| GAG biosynthesis (all) | *Xylt1* |
| | *Xylt2* |
| | *B4galt7* |

| | |
|---|---|
| | *B3galt6* |
| | *B3gat3* |
| GAG biosynthesis (HS) | *Ext1* |
| | *Ext2* |
| | *Extl1* |
| | *Extl2* |
| | *Extl3* |
| GAG biosynthesis (CS) | *Csgalnact1* |
| | *Csgalnact2* |
| | *Chpf* |
| | *Chsy1* |
| | *Chsy3* |
| HA synthesis | *Has1* |
| | *Has2* |
| | *Has3* |
| HS sulfation | *Glce* |
| | *Hs6st1* |
| | *Hs6st2* |
| | *Hs6st3* |
| | *Ndst1* |
| | *Ndst2* |
| | *Ndst3* |
| | *Ndst4* |
| | *Hs3st1* |
| | *Hs3st2* |
| | *Hs3st3A* |
| | *Hs3st3B* |
| | *Hs3st4* |
| | *Hs3st5* |
| | *Hs3st6* |
| | *Hs2st1* |
| CS sulfation | *Chst3* |

| | |
|---|---|
| | *Chst11* |
| | *Chst12* |
| | *Chst13* |
| | *Chst14* |
| | *Chst15* |
| | *Ust* |

**Enzyme-linked immunosorbent assay (ELISA)**

Circulating HS concentrations were quantified in mouse serum using a commercially available enzyme-linked immunosorbent assay (ELISA) (Finetest) according to the manufacturer's instructions. Whole blood was collected from femoral artery at the experimental endpoint and serum collected and stored at -80°C until analysis. Serum samples were thawed only once prior to ELISA to minimise degradation. Optical density values were measured at 450 nm and serum heparan sulphate concentrations were interpolated from a regression line. Quality control samples and blank wells were included on each plate where appropriate. Duplicate measurements were averaged to generate a single value for each biological replicate.  All samples from an individual experiment were analysed on the same ELISA plate wherever possible to minimise inter-assay variability.

**Quantitative real-time polymerase chain reaction (qPCR)**

Lung tissue was collected into 500 µL RNAlater Stabilization Solution (Thermo Fisher Scientific) immediately following tissue collection and stored overnight at 4°C before long-term storage at -80°C. Prior to RNA extraction, samples were thawed on ice and total RNA was isolated from approximately 2 mg of lung tissue using the RNeasy Mini Kit (Qiagen) according to the manufacturer's instructions. RNA concentration and purity were assessed using a NanoDrop spectrophotometer.

Complementary DNA (cDNA) was synthesised from purified RNA using SuperScript Reverse Transcriptase (Thermo Fisher Scientific) in combination with Oligo(dT)12–18 primers (Thermo Fisher Scientific), according to the manufacturer's instructions. Quantitative real-time PCR was performed to determine relative expression of Ext1, Extl2, Extl3, B4galt7, Mmp9 and Hpse using the primer sequences listed in

Supplementary Table 5. Reactions were prepared using SYBR Green Master Mix (Thermo Fisher Scientific) and analysed using a QuantStudio 12K Flex Real-Time PCR System (Thermo Fisher Scientific).

Five serial 1:4 dilutions of pooled cDNA were included to generate standard curves and assess amplification efficiency. Gene expression was quantified using the relative standard-curve method and normalised to expression of the housekeeping gene *Hprt*. Relative expression values are presented relative to *Hprt*.

**Supplementary Table 5. Primer sequences used to characterise expression of glycocalyx-modifying enzymes in the lung**

| **Gene** | **Sequence** | |
|---|---|---|
| *Ext1* | Fwd | 5’-GGAGTTGCCATTCTCCGA-‘3 |
| | Rev | 5’-TAAGCCTCCCACAAGAACTG-‘3 |
| *Extl2* | Fwd | 5’-AACCGACTCCAGGTCTT-‘3 |
| | Rev | 5’-AACTGTAGATCCCGGAGG-‘3 |
| *Extl3* | Fwd | 5’-CAATTCCAACTACTCCTGTG-‘3 |
| | Rev | 5’-GGGCATCACATAAGAATACAG-‘3 |
| *B4Galt7* | Fwd | 5’-AGAGGATGATGAATTCTACCG–‘3 |
| | Rev | 5’-AACTGTTCCTGTTTTTGAGC–‘3 |
| *Hpse* | Fwd | 5’-GAGAGAGCCAGTAATCAGGTAAAG-‘3 |
| | Rev | 5’-GGGCATAGAAGTCGTGATGAG-‘3 |
| *Mmp9* | Fwd | 5’-CTTCCAGTACCAAGACAAAG-‘3 |
| | Rev | 5’-ACCTTGTTCACCTCATTTTG-‘3 |
| *Hprt* | Fwd | 5’-TCCTCCTCAGACCGCTTTT-‘3 |
| | Rev | 5’-CCTGGTTCATCATCGCTAATC-‘3 |

**Statistical Analysis**

Statistical analyses were performed using GraphPad Prism version 10 (GraphPad Software, San Diego, CA, USA). Comparisons between two independent groups, including experiments comparing PBS control and IAV-infected mice at a single 24 h time point, were analysed using an unpaired two-tailed Student's t-test. Influenza infection time-course experiments containing multiple time points were analysed using one-way ANOVA followed by Dunnett's multiple-comparisons test, with each post-infection time point compared with the PBS control group.

Experiments containing both genotype (WT versus KO) and treatment (PBS versus IAV) were analysed using two-way ANOVA. The main effect of genotype was used to assess overall differences between WT and KO mice across treatment groups. Where individual group comparisons were performed, Sídak's multiple-comparisons test was applied. Data are presented as mean ± SD or SEM as specified in the corresponding figure legends. A $P$ value $< 0.05$ was considered statistically significant.

### Supplementary Figures:

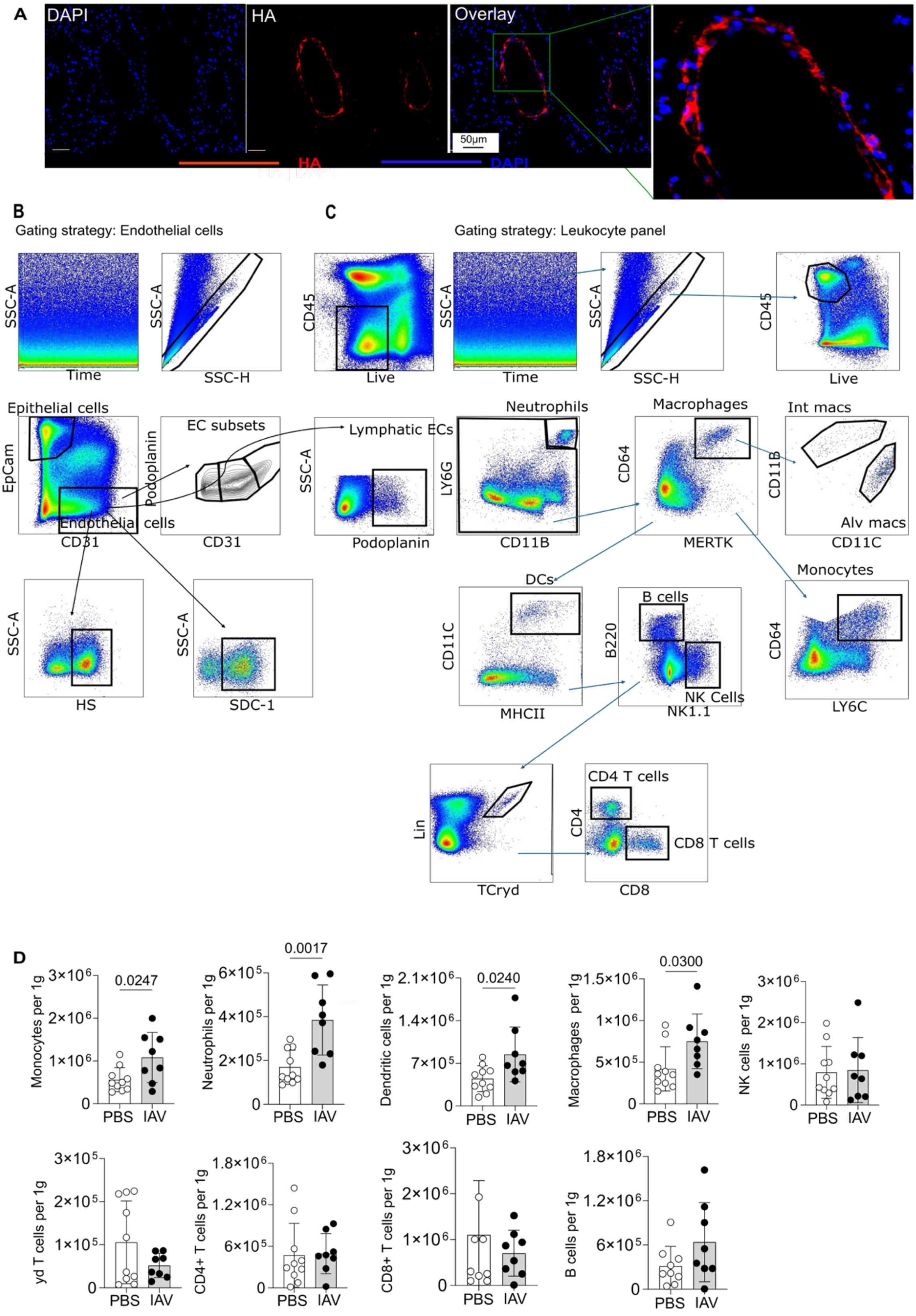

**Fig. S1. Endothelial and immune-cell phenotyping 24 h following IAV infection.** (A) Representative immunofluorescence images of uninfected mouse lung sections stained for nuclei (DAPI) in blue and hyaluronan (HA) in red. HA was detected using hyaluronan-binding protein (HBP) followed by fluorophore-conjugated streptavidin. (scale bar = 50 μm). (B) Representative gating strategy for flow-cytometric identification of mouse lung endothelial cells. (C) Representative gating strategy for flow-cytometric identification of mouse lung immune-cell populations. (D) Quantification of individual immune-cell populations in the lungs of mice 24 h following PBS control or IAV infection. Each point represents one mouse; n = (8-10) mice per group. D was analysed by unpaired t-test with p values indicated.

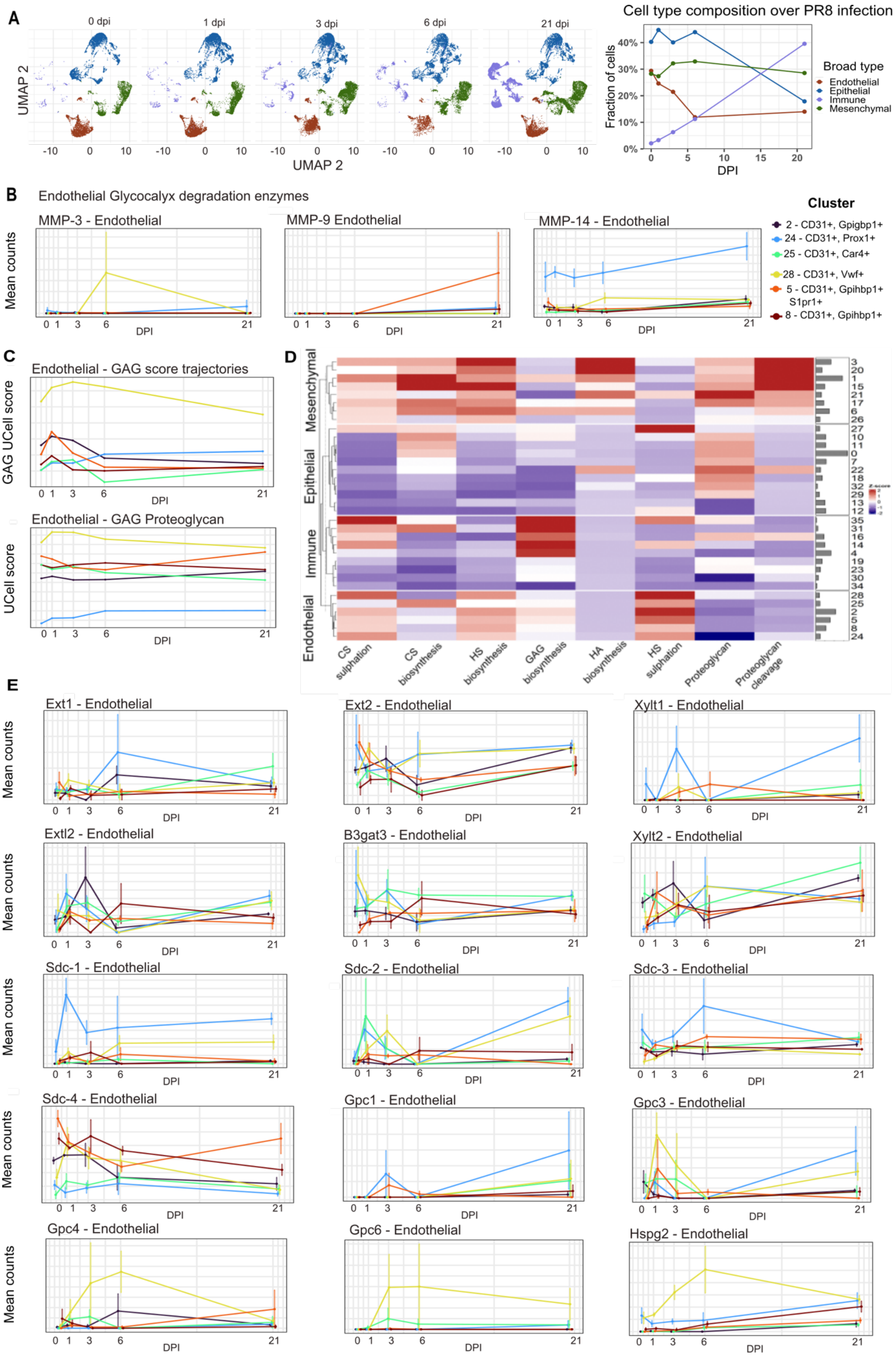

A
0 dpi
1 dpi
3 dpi
6 dpi
21 dpi
UMAP 2
UMAP 2
Cell type composition over PR8 infection
Fraction of cells
DPI
Broad type
Endothelial
Epithelial
Immune
Mesenchymal
B
Endothelial Glycocalyx degradation enzymes
MMP-3 - Endothelial
MMP-9 Endothelial
MMP-14 - Endothelial
Mean counts
DPI
Cluster
2 - CD31+, Gpigbp1+
24 - CD31+, Prox1+
25 - CD31+, Car4+
28 - CD31+, Vwf+
5 - CD31+, Gpihbp1+ S1pr1+
8 - CD31+, Gpihbp1+
C
Endothelial - GAG score trajectories
GAG UCell score
Endothelial - GAG Proteoglycan
UCell score
D
Mesenchymal
Epithelial
Immune
Endothelial
CS sulphation
CS biosynthesis
HS biosynthesis
GAG biosynthesis
HA biosynthesis
HS sulphation
Proteoglycan
Proteoglycan cleavage
E
Ext1 - Endothelial
Ext2 - Endothelial
Xylt1 - Endothelial
Extl2 - Endothelial
B3gat3 - Endothelial
Xylt2 - Endothelial
Sdc-1 - Endothelial
Sdc-2 - Endothelial
Sdc-3 - Endothelial
Sdc-4 - Endothelial
Gpc1 - Endothelial
Gpc3 - Endothelial
Gpc4 - Endothelial
Gpc6 - Endothelial
Hspg2 - Endothelial

**Fig. S2. The endothelial glycocalyx transcriptional response during the influenza time course.** (A) uMAP clustering and fractional analysis of endothelial, epithelial, immune and mesenchymal cells in the mouse lung at the indicated time points following IAV infection. (B) Analysis of *MMP 3, 9 and 14* gene expression in different endothelial cell subsets across the IAV infection time course. (C) Analysis of grouped genes categorised as relating to GAG production of proteoglycan production in different endothelial cell subsets across the influenza infection time course. (D) Heatmap analysis of different GAG gene families in all cell types. (E) Gene trajectory analysis of indicated individual genes in different endothelial cell subsets across the IAV infection time course.

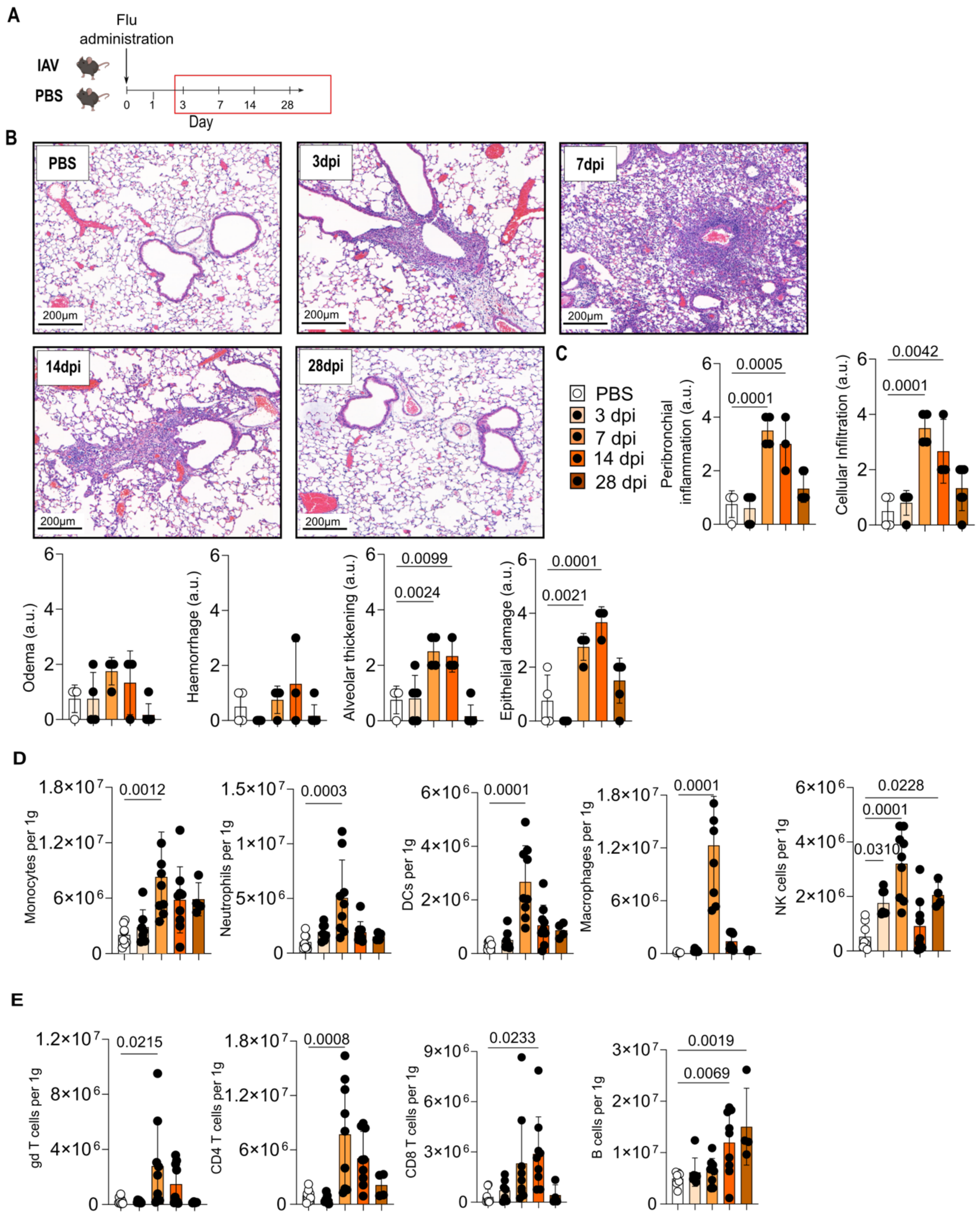

A
Flu administration
IAV
PBS
0 1 3 7 14 28
Day
B
PBS
3dpi
7dpi
14dpi
28dpi
200μm
C
PBS
3 dpi
7 dpi
14 dpi
28 dpi
Peribronchial inflammation (a.u.)
0.0005
0.0001
Cellular Infiltration (a.u.)
0.0042
0.0001
Odema (a.u.)
Haemorrhage (a.u.)
Alveolar thickening (a.u.)
0.0099
0.0024
Epithelial damage (a.u.)
0.0001
0.0021
D
Monocytes per 1g
0.0012
Neutrophils per 1g
0.0003
DCs per 1g
0.0001
Macrophages per 1g
0.0001
NK cells per 1g
0.0228
0.0001
0.0310
E
gd T cells per 1g
0.0215
CD4 T cells per 1g
0.0008
CD8 T cells per 1g
0.0233
B cells per 1g
0.0019
0.0069

**Fig. S3. Characterising a time-course model of the mouse lung influenza immune response.** (A) Schematic of the longitudinal mouse IAV infection model. (B) Haematoxylin and eosin staining of mouse lung tissue section from across the IAV infection time course. (C) Quantification of lung tissue damage scores across the IAV infection time course. Each point represents one mouse (n = 3-5) mice per group. (D) Quantification of immune cell populations within the mouse lung across the influenza infection time course expressed as cells per gram of tissue.  Each point represents one mouse; n = (8-10). C-E were analysed by one-way ANOVA with p values indicated.

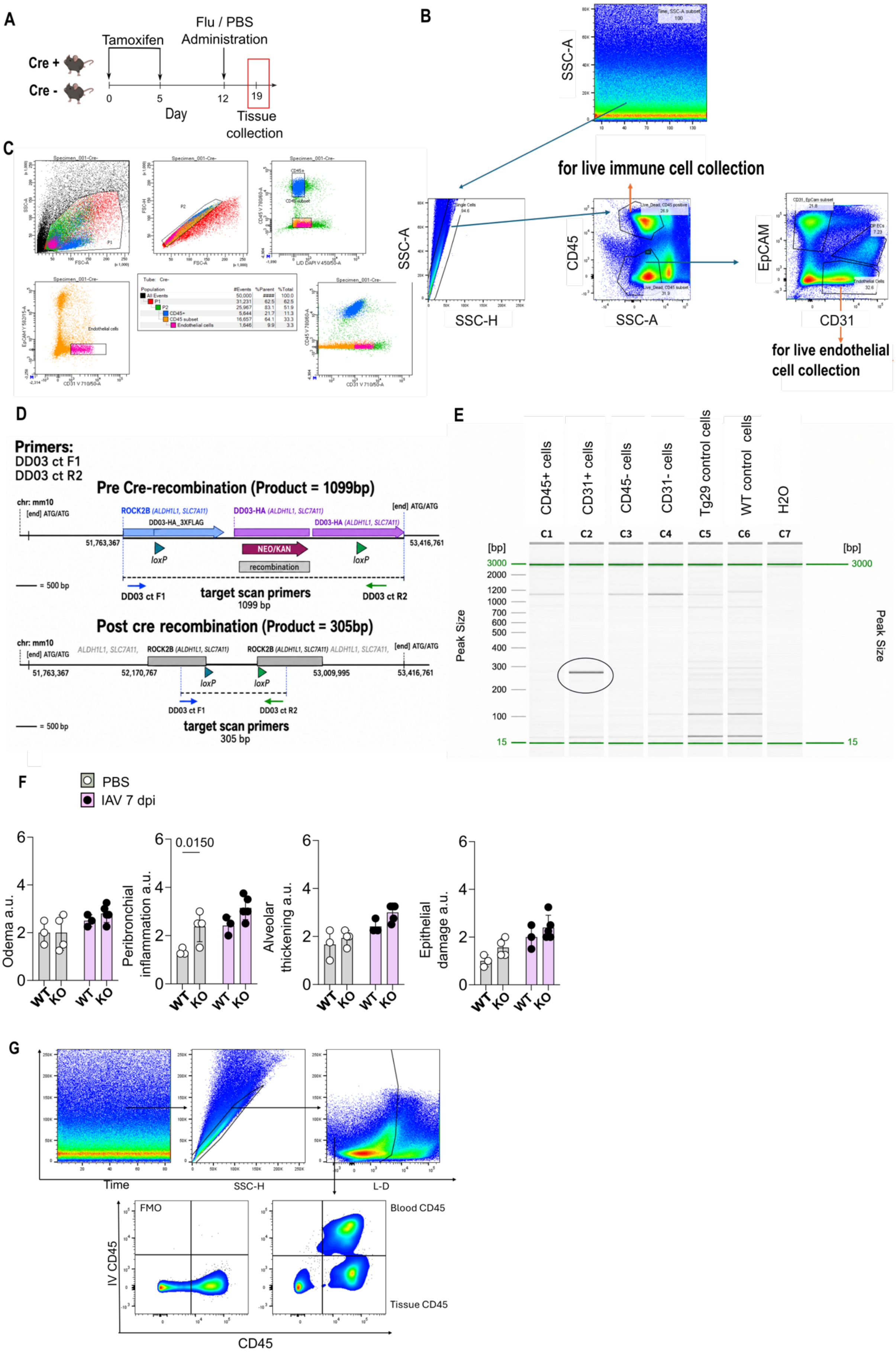
A
Flu / PBS Administration
Tamoxifen
Cre +
Cre -
0
5
12
19
Day
Tissue collection
B
SSC-A
for live immune cell collection
SSC-A
SSC-H
CD45
SSC-A
EpCAM
CD31
for live endothelial cell collection
C
D
Primers:
DD03 ct F1
DD03 ct R2
Pre Cre-recombination (Product = 1099bp)
NEO/KAN
recombination
loxP
target scan primers
1099 bp
Post cre recombination (Product = 305bp)
target scan primers
305 bp
E
CD45+ cells
CD31+ cells
CD45- cells
CD31- cells
Tg29 control cells
WT control cells
H2O
C1
C2
C3
C4
C5
C6
C7
[bp]
Peak Size
F
PBS
IAV 7 dpi
Odema a.u.
Peribronchial inflammation a.u.
0.0150
Alveolar thickening a.u.
Epithelial damage a.u.
WT
KO
G
Time
SSC-H
L-D
FMO
Blood CD45
Tissue CD45
IV CD45
CD45

**Fig. S4. Developing a new cell-specific glycocalyx KO mouse model and characterising its' response to influenza infection.** (A) Schematic of the experimental design and tamoxifen induction regimen used to induce endothelial cell-specific Cre-mediated recombination prior to IAV infection and lung tissue collection. (B) Representative flow cytometry gating strategy used to identify pulmonary $CD45^-$ $CD31^+$ endothelial cells and $CD45^+$ immune cells from naive lung tissue for endothelial cell and immune cell sorting. Live single cells were gated followed by identification of endothelial cells as $CD45^-CD31^+EpCAM^-$ populations and identification of immune cells as $CD45^+$ cell populations. (C) Representative flow cytometry plots showing cell sorting strategy for endothelial and $CD45^+$ populations used for downstream validation analyses. (D) Schematic representation of the *B4galt7* locus showing primer locations and expected PCR products before and after Cre-mediated recombination. (E) Representative PCR gel demonstrating recombination of the *B4galt7* allele within endothelial cells (circled lane) with absence of recombination in $CD45^+$ immune cell populations. (F) Mouse lung histological tissue damage scoring across the IAV infection time course in WT (*Cdh5*-Cre/ERT2$^{-/-}$ x *B4Galt7*$^{fl/fl}$) and endothelial GAG KO (*Cdh5*-Cre/ERT2$^{+/-}$ x *B4Galt7*$^{fl/fl}$) mice. Each point represents one mouse; n = (3-5) mice per group. (G) Representative flow cytometry plots showing gating strategy illustrating intravenous anti-CD45 (ivCD45) labelling for separation of tissue and vascular localised immune cells.

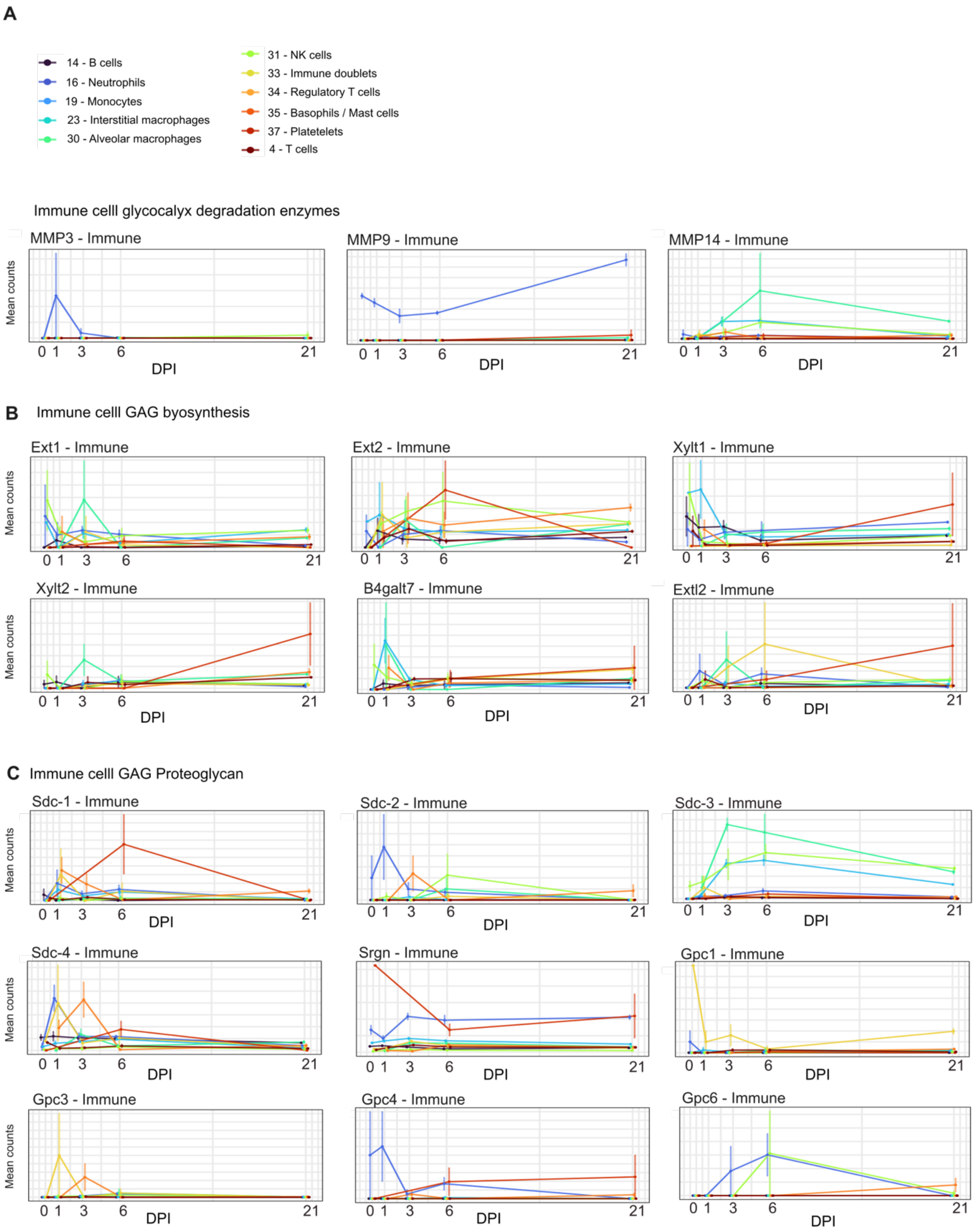
A
14 - B cells
16 - Neutrophils
19 - Monocytes
23 - Interstitial macrophages
30 - Alveolar macrophages
31 - NK cells
33 - Immune doublets
34 - Regulatory T cells
35 - Basophils / Mast cells
37 - Platetelets
4 - T cells
Immune celll glycocalyx degradation enzymes
MMP3 - Immune
MMP9 - Immune
MMP14 - Immune
Mean counts
0 1 3 6 21
DPI
B Immune celll GAG byosynthesis
Ext1 - Immune
Ext2 - Immune
Xylt1 - Immune
Xylt2 - Immune
B4galt7 - Immune
Extl2 - Immune
C Immune celll GAG Proteoglycan
Sdc-1 - Immune
Sdc-2 - Immune
Sdc-3 - Immune
Sdc-4 - Immune
Srgn - Immune
Gpc1 - Immune
Gpc3 - Immune
Gpc4 - Immune
Gpc6 - Immune

**Fig. S5. The immune cell glycocalyx transcriptional response during the IAV infection time course.** Longitudinal trajectory analysis of specific immune cell GAG synthesis and remodelling gene expression across the IAV infection time course. (A) Glycocalyx degradation enzymes, (B) GAG biosynthesis genes and (C) proteoglycan core genes.

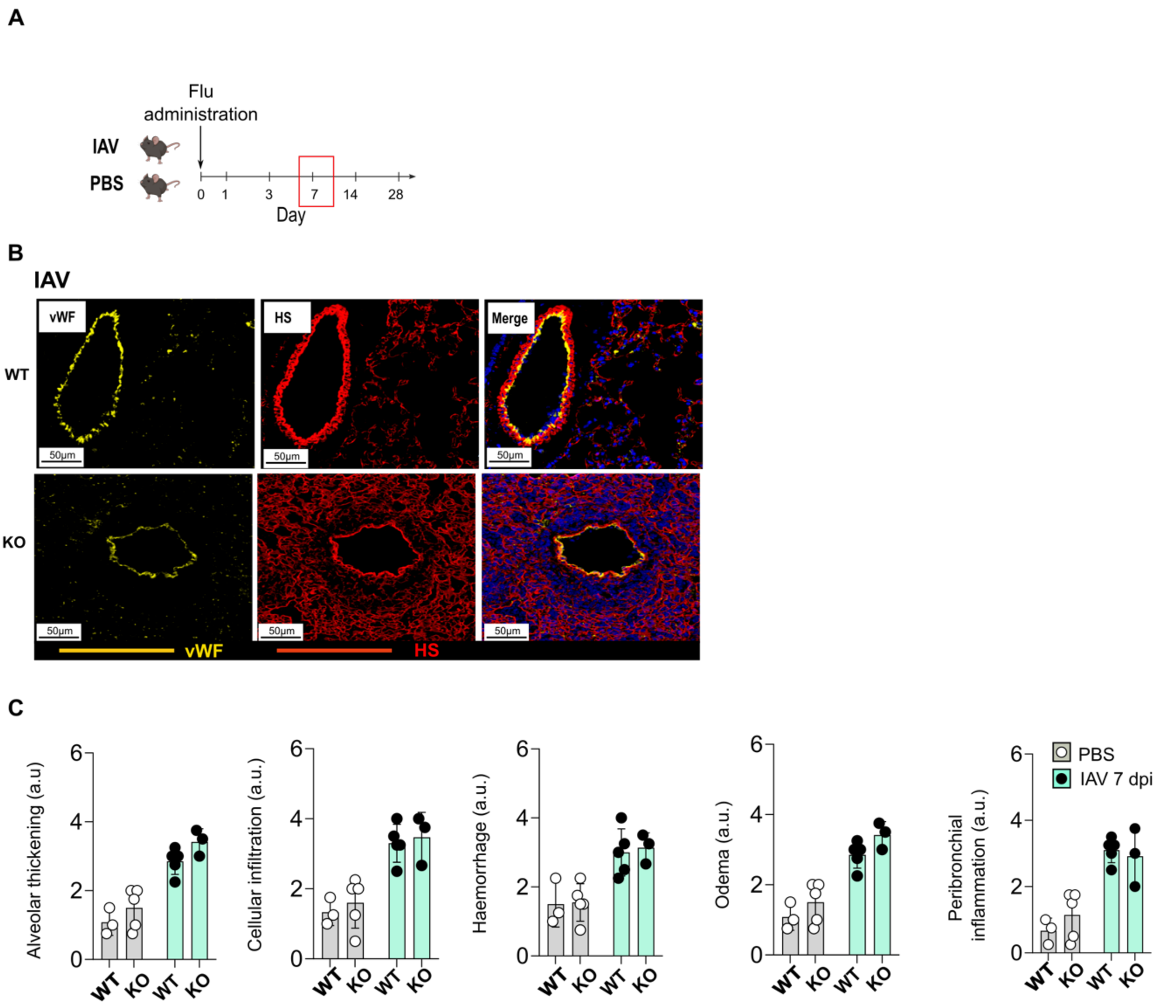
A
Flu administration
IAV
PBS
0 1 3 7 14 28
Day
B
IAV
vWF
HS
Merge
WT
KO
50µm
vWF
HS
C
Alveolar thickening (a.u)
Cellular infiltration (a.u.)
Haemorrhage (a.u.)
Odema (a.u.)
Peribronchial inflammation (a.u.)
WT
KO
PBS
IAV 7 dpi

**Fig. S6. Characterising the murine IAV immune response in the *Hpse* KO mouse.** (A). Schematic of IAV infection and tissue collection. (B) Immunofluorescence analysis of vascular ($vWF^+$) heparan sulphate (HS) in lung tissue from wild-type (WT) and *Hpse* KO mice 7 days following PBS control or IAV infection. (C) Blinded histopathological assessment of individual lung tissue damage parameters in WT and *Hpse* KO mice 7 days following PBS control or IAV infection. Each point represents one mouse; n = (3-5) mice per group.
